# Association of *CDH11* with ASD revealed by matched-gene co-expression analysis and mouse behavioral studies

**DOI:** 10.1101/2020.02.04.931121

**Authors:** Nan Wu, Yue Wang, Jing-Yan Jia, Yi-Hsuan Pan, Xiao-Bing Yuan

## Abstract

A large number of putative risk genes of autism spectrum disorder (ASD) have been reported. The functions of most of these susceptibility genes in developing brains remain unknown, and a causal relationship between their variations and autism traits has not been established. The aim of this study is to predict putative risk genes at the whole-genome level based on the analysis of gene co-expression with a group of high confidence ASD risk genes (hcASDs). Results showed that three gene features, including gene size, mRNA abundance, and guanine-cytosine content, affect genome-wide co-expression profiles of hcASDs. To circumvent the interference of these gene features on gene co-expression analysis (GCA), we developed a method to determine whether a gene is significantly co-expressed with hcASDs by statistically comparing the co-expression profile of this gene with hcASDs to that of this gene with permuted gene sets of feature-matched genes. This method is referred to as “matched-gene co-expression analysis” (MGCA). With MGCA, we demonstrated the convergence in developmental expression profiles of hcASDs and improved the efficacy of risk gene prediction. Results of analysis of two recently reported ASD candidate genes, *CDH11* and *CDH9,* suggested the involvement of *CDH11*, but not *CDH9*, in ASD. Consistent with this prediction, behavioral studies showed that *Cdh11*-null mice, but not *Cdh9*-null mice, have multiple autism-like behavioral alterations. This study highlighted the power of MGCA in revealing ASD-associated genes and the potential role of CDH11 in ASD.

## Introduction

Autism spectrum disorder (ASD) is a heterogeneous neurodevelopmental condition with a complex genetic basis ^[1, 2]^. A large number of putative risk genes have been identified by genetic linkage analyses, genome-wide association studies (GWAS), whole-exome sequencing (WES), or whole-genome sequencing (WGS) ^[3–5]^. However, the functions of most of these putative risk genes in developing brains remain unknown. For some novel risk genes, the genetic evidence supporting their association with ASD is not sufficient. Therefore, a causal relationship between the variations of many risk genes and autism traits has not been established. In order to prioritize the investigation of genes and signaling pathways of high relevance to ASD, a method to determine the functional importance of a large group of putative risk genes is vital.

The highly diverse ASD risk genes are believed to functionally converge in several common molecular pathways closely related to ASD, such as the Wnt signaling pathway, the mammalian target of rapamycin (mTOR) pathway, and dendrite development and synaptic remodeling pathways ^[3, 6]^. Consistent with the functional convergence of ASD risk genes, several studies showed the convergence of developmental expression profiles of a large group of risk genes ^[7, 8]^. It is generally believed that genes with similar expression profiles are co-regulated or have related functions ^[7, 9]^. The co-expression of genes within a biological pathway is a strong indication of their shared functions ^[9]^. Based on this concept, computational analyses of various brain transcriptomes have been conducted to identify potential co-expression networks of ASD risk genes and to discover brain circuits that may be affected by risk genes ^[7, 8, 10, 11]^. In these studies, the correlation coefficient (CC) of a pair of genes is calculated based on their expression levels in different brain regions or developmental stages. Genome-wide gene co-expression networks are constructed by setting an empirically determined threshold of CC ^[7]^. Genetic mutation and protein interaction data have also been incorporated into gene co-expression analysis (GCA) along with novel data analysis algorithms, such as machine learning, to predict putative ASD risk genes and their convergent molecular pathways^[12]^. A major limitation in most of these studies is the lack of ways to overcome the potential effects of confounding factors such as the size, expression level (mRNA abundance), and guanine-cytosine (GC) content of genes on the result of GCA ^[13]^. Most ASD risk genes are large genes with a higher expression level in the brain than in other tissues ^[14]^. It is unclear whether the size or expression level of an ASD gene affects its co-expression with other genes. It is also unclear whether the convergent pattern of developmental expression profiles is specific to ASD risk genes or the common property of genes with similar features, such as large gene size and high mRNA abundance ^[13]^.

In this study, we discovered that three gene features, including mRNA abundance, gene size (genomic DNA length), and GC content of a gene, affect gene co-expression profiles in the brain. We developed a novel method called “matched-gene co-expression analysis” (MGCA) (**Fig. 1**) to examine whether a gene exhibits significant co-expression with a group of high-confidence ASD risk genes (hcASDs) independent of confounding gene features. This was accomplished by statistically comparing the co-expression level of a gene with the hcASD gene set to that of this gene with a large number of permuted gene sets of matched features. MGCA effectively predicted putative ASD-associated genes and revealed functionally important convergent molecular pathways of these genes. MGCA also revealed “homophilic cell adhesion” as one of the most significantly converged pathways of risk genes. Results of further analysis of *CDH11* and *CDH9*, two ASD candidate genes belonging to the cadherin family of adhesion molecules, suggested the involvement of *CDH11*, but not *CDH9*, in ASD. This prediction was supported by mouse behavioral studies using *Cdh11-* and *Cdh9*-null mice.

**Figure 1.**
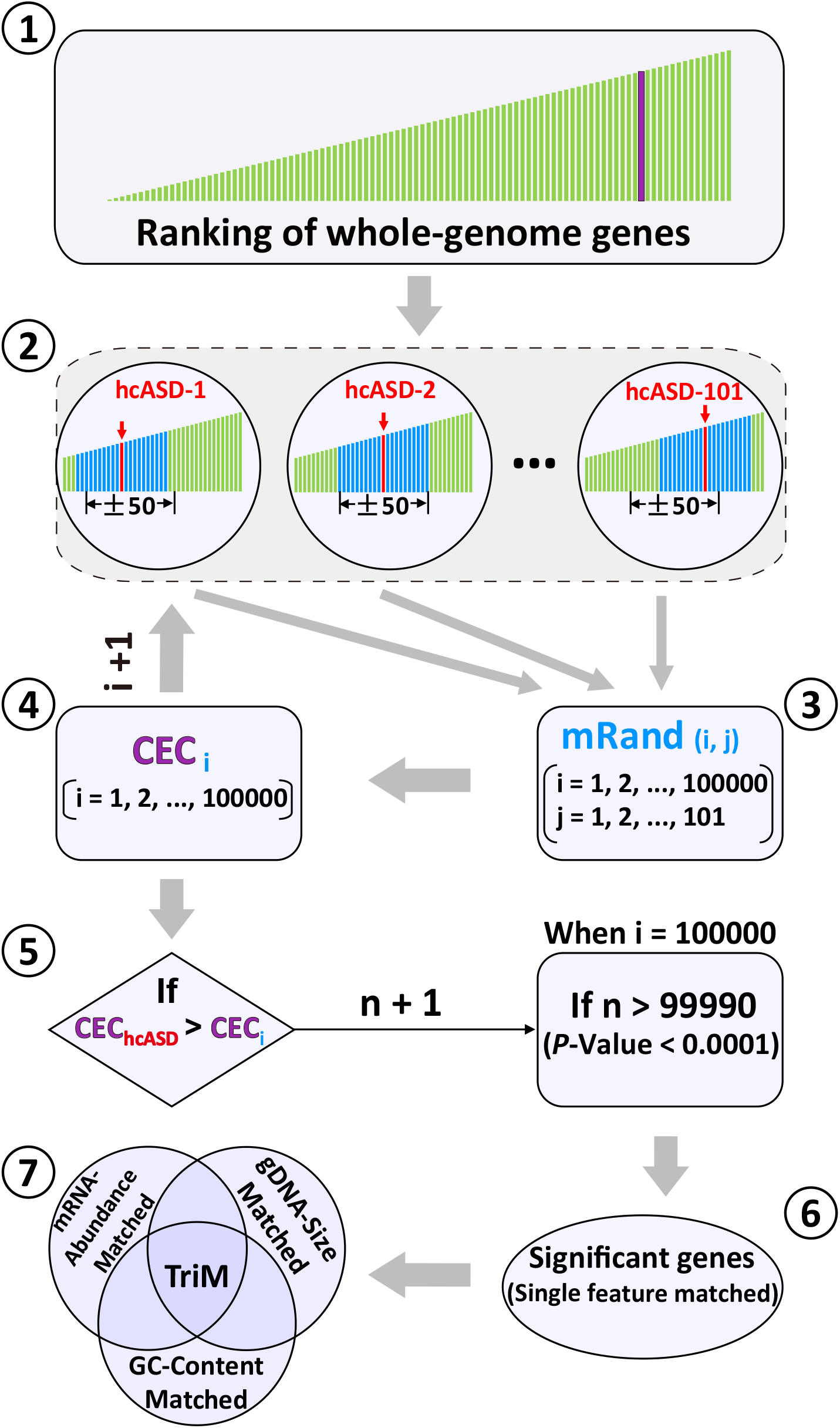
Flowchart of the matched gene co-expression analysis (MGCA). Whole-genome genes were ranked by gene features, creating various gene lists ①. For each gene in the hcASD gene set (hcASD-1, hcASD-2, ··· to hcASD-101), a feature-matched gene was randomly selected from the range of 50 genes above and 50 genes below (±50 range) this gene in the ranked gene lists ② to generate a matched random gene set (mRand) ③. For each gene in the whole genome, its co-expression coefficient (CEC) with each of the mRand gene set (CEC_i_) was computed ④, and 100,000 permutations were conducted (i=1,2,…100000). Each CEC_i_ was compared to the CEC of the same gene with the hcASD gene set (CEC_hcASD_). Count the number of permutations with CEC_i_ < CEC_hcASD_ (n started from 0 for each gene evaluated, n+1 if CEC_i_ < CEC_hcASD_) ⑤. At the end of 100000 permutations, genes with n > 99990, which corresponded to a permutation *P*-value < 0.0001, were considered significantly co-expressed with hcASDs under a feature-matched condition ⑥. Genes significantly co-expressed with hcASDs under all three feature-matched conditions were defined as Triple-matched genes (TriM) ⑦.

## Materials and Methods

### Data filtering and computation of co-expression coefficient

The human brain transcriptome dataset from BrainSpan (www.brainspan.org) (RNA-Seq Gencode v10) was used for GCA. This dataset contained 256 transcriptomes of 16 different brain regions. The developmental stages ranged from post-conception week 8 (PCW8) to 40 years old (40Y). Normalized mRNA expression values were represented by RPKM (Reads Per Kilobase Per Million Mapped Reads). The average mRNA expression level of each gene in all tissues was considered as the mRNA abundance level of a gene. Gene lengths were determined based on gene annotations provided by the National Center for Biotechnology Information (NCBI). The GC content of a gene was obtained from Ensembl Genome Browser. Based on statistical analyses of genetic data described previously, 101 risk genes that reached a genome-wide significance threshold (false discovery rate, FDR ≤ 0.1) ^[15]^ were used as the hcASD gene set (high confidence ASD risk gene set,**Table S1**). Genes with an abundance level lower than the lowest abundance level of hcASDs were filtered out (**Table S1**). Perl scripts were written to conduct most calculations. Pair-wise Pearson’s correlation coefficient (CC) was used to indicate the tendency of co-expression of a gene pair. Heatmaps were constructed with the software R based on the CC matrix of 1/100 evenly distributed genes. The mean CC was defined as co-expression coefficient (CEC), which indicates the tendency of co-expression of a gene with a specific set of genes (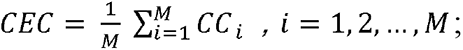 where *M* indicates the total gene number of a gene set) or the tendency of co-expression of two gene sets (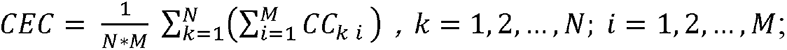 where *M* and *N* represent the total gene number of two different gene sets, respectively).

### Gene set definition

After data filtering, a total of 12,250 genes with information on gene length, mRNA level, and GC content available were identified and used for the study (**Table S1**). In addition to the hcASD gene set, the following gene sets were also used: cASD (combined ASD genes), mRand (matched random genes), Rand (random genes), TriM (triple-matched genes), Top (Top-ranked genes by CEC value), TriM-only (genes only in TriM gene set), and Top-only (genes only in Top gene set). The cASD gene set is a combined set of ASD-associated genes containing 514 non-redundant genes from nine different sets of previously reported ASD-associated genes (**Table S5**)^[7, 15–21]^. Each mRand gene set contained 101 genes, with one of the three gene features of each gene matched with that of the corresponding hcASD gene in the hcASD gene set. To generate the mRand gene sets, each feature-matched gene was randomly selected within the ±50 range of the corresponding hcASD gene in the ranked gene list using a Perl Script. Each Rand gene set contained 101 genes randomly selected from the whole gene list using a Perl Script without considering matched gene features. TriM was the set of MGCA-revealed genes that exhibited significant co-expression with the hcASD gene set under all three matched conditions (see **Figs. 1, 4A**). The Top gene set contained top-ranked genes with the highest CEC values with hcASD. TriM-only and Top-only gene sets contained non-overlapped genes present only in the TriM or the Top gene set.

### Gene ontology (GO) analysis

GO analysis was performed using DAVID v6.8 (http://david.ncifcrf.gov/tools.jsp), and the human whole-genome genes provided by DAVID were used as the background list. A corrected *P*-value of 0.05 (Benjamini-Hochberg method) was used.

### Pathway enrichment analysis

Metascape (http://metascape.org) was used to perform pathway enrichment analysis and draw heatmaps. Pairwise similarities between any two significant terms were computed based on Kappa-test scores. The enriched terms were then hierarchically clustered into a tree with a kappa score of 0.3 as the threshold. Boxes were colored according to their *P*-values. Gray boxes indicate a lack of enrichment for a specific GO term.

### Ethics approval

Animal care and handling were performed according to the guidelines for the Care and Use of Laboratory Animals of the National Institutes of Health. All animal experiments were approved by the Animal Care and Use Committees of Hussman Institute for Autism (06012015D), University of Maryland School of Medicine (0515017), and East China Normal University (m20190236).

### Animals

*Cdh9*-null mice (*Cdh9*^Laz^, C57BL/6-ICR mixed background) ^[22]^ were provided by Dr. Joshua R. Sanes at Harvard University. *Cdh11*-null mice ^[23]^ were from the Jackson Labs (*Cdh11^tm1Mta^*/HensJ, https://www.jax.org/strain/023494, C57BL/6-129Sv-CD-1 mixed background). All mice were housed in groups of five with free access to food and water and kept on a 12-hour light/dark cycle. Mice were tattooed on the tail using fluorescent ink for identification. A UV flashlight was used to visualize the tattooed identification numbers. All behavioral tests were conducted during the daytime on mice 2-5 months of age. The experimenter was blind to the genotype of the animal during behavioral experiments. The surface of the apparatus for behavioral tests was cleaned with 50% ethanol between tests. At least 5□ min between cleaning and the next test was allowed for ethanol evaporation and odor dissipation.

### Genotyping

Genotyping of *Cdh9*-null mice was done by PCR as previously described ^[22]^. The PCR product for the wildtype (WT) *Cdh9* allele was 550 bp amplified with the primer pair *Cdh9*-P1 (CCA CTA CAG GAA ACC TTT GGG TT) and *Cdh9*-P3 (ATG CAA ACC ATC AGG TAT ACC AAC C), and that of the mutant allele was 430 bp amplified with the primer pair *Cdh9*-P1 and *Cdh9*-P2 (CGT GGT ATC GTT ATG CGC CT). The annealing temperature for *Cdh9* PCRs was 63°C. For genotyping of *Cdh11*-null mice, the primer pair *Cdh11*-P1 (CGC CTT CTT GAC GAG TTC) and *Cdh11*-P2 (CAC CAT AAT TTG CCA GCT CA) were used for amplification of the mutant allele, and the primer pair *Cdh11*-P3 (GTT CAG TCG GCA GAA GCA G) and *Cdh11*-P2 were used for the WT allele. The annealing temperatures for PCR were 63.1°C and 56°C for the mutant and WT alleles, respectively. The sizes of the PCR products for the mutant and WT alleles were 500 bp and 400 bp, respectively.

### Behavioral tests

Mice of 3-5 months old were used for these behavioral tests. Animals were handled before the test (10 min/day for 3 days). The general order of behavioral tests was the open field test, elevated plus maze, sociability test, rotarod test, and gripping force test. Animals rested for at least 3 days after finishing one behavioral experiment. During all behavioral experiments, the experimenter was blind to mouse genotypes. Three batches of mice were analyzed, and data were pooled for analysis.

#### Open field test

The test mouse was allowed to freely explore the open field arena (50 cm × 50 cm) for 30 min. The mouse’s motion was videoed and tracked by an automated tracking system (EthoVision XT 11.5), which also recorded rearing, hopping, turning, self-grooming, moving time, total moving distance, and time spent in the center of the arena (1/2 of total size).

#### Elevated plus maze test

The standard elevated plus maze (EPM) apparatus consisted of two open and two closed arms, 30 × 5 cm each, connected by a central platform (5 cm × 5 cm). The maze was 30 cm off the ground. The test mouse was gently placed on the central platform with its head facing one closed arm and was allowed to explore for 10 min freely. The mouse’s time stayed in the two open arms and the frequency of open arm entry were recorded.

#### Grip strength test

The test mouse was placed on a metal grid on top of a transparent chamber. The grid was quickly inverted, and the time for the mouse to drop off the grid was determined. Five consecutive trials were carried out, and the average hanging time for each mouse was calculated. The maximum hanging time was set for 1 min. After 1 min of hanging, the trial was stopped, and the hanging time was recorded as 1 min.

#### Horizontal bar test

The mouse was gently placed on a metal wire, with the two forepaws gripping the wire. The length of time which the mouse hung on the wire was measured. The maximum hanging time was set for 1 min. The average hanging time was calculated from 5 consecutive trials.

#### Rotarod test

Mice were habituated to the rotarod apparatus (Harvard Apparatus 760770) by leaving them on the low-speed rotating rod (4 rpm) for 5 min each day for 3 days and tested on the fourth day on the accelerating rod. The time and the maximum rotation speed that the test mouse maintained the balance on the rotating rod were measured. Five consecutive trials were done for each mouse.

#### Social preference test

A modified three-chamber apparatus was used. The apparatus comprised 3 rectangular (25 cm × 38 cm) chambers made of white Plexiglas with a 13 cm gate connecting the two side chambers to the middle chamber. A 3-sided (13 cm wide for each side) fence made of transparent Plexiglas was placed inside each side chamber facing the door of the side chambers, creating a 13 cm × 13 cm square area separated from the side chambers but connected to the middle chamber through the door (**Fig. 7A**). The two side chambers were covered by transparent Plexiglas to minimize odorant diffusion. The test mouse was placed inside the middle chamber and freely explored the middle chamber and the square zone in each side chamber for 10 min. Three social partner mice were then placed into the fenced area in one side chamber, and the test mouse was allowed to explore for another 10 min freely. Another 3 social partner mice were then placed in the other side chamber, and the behavior of the test mouse was tracked for 10 min. The time that the test mouse spent in each chamber was measured.

### Experimental Design and Statistical Analysis

The effects of three different gene features on gene co-expression profiles were first analyzed. MGCA (**Fig. 1**) was then performed to determine the convergence in developmental expression profiles of hcASDs and detect genes that were significantly co-expressed with hcASDs in the whole genome. The effectiveness of MGCA in predicting putative ASD-associated genes was analyzed by comparing its results with those of GCA, which does not consider. *CDH11* and *CDH9* were selected as example genes, and mouse behavioral experiments were conducted in gene knockout mice to verify the findings of MGCA.

Data are presented as mean ± standard error of the mean (SEM). The upper fence test and Grubbs’ test were performed to evaluate whether the hcASD-hcASD expression level (CEC value) was significantly higher than the CEC values of feature-matched (mRand-mRand; hcASD-mRand) or non-matched non-hcASD gene sets (Rand-Rand; hcASD-Rand). Grubb’s test was done using the “grubbs.test” script in the R software package. The false discovery rate (FDR) of a gene was determined by the frequency of this gene significantly co-expressed (*P* < 0.001) with 5,000 mRand gene sets determined by MGCA. Gene enrichment *P*-value was determined with the Chi-Square test. Pathway enrichment *P*-values were determined with Metascape. Behavioral data were analyzed by Student *t*-test and by one-way ANOVA followed by Dunnett’s *t*-test as post hoc analysis using SPSS (IBM, Armonk, USA) or GraphPad Prism (GraphPad Software, La Jolla, CA, USA).

### Availability of data and materials

Perl scripts for data analysis are available on GitHub (https://github.com/wunan124/MGCA)

## Results

### Effects of gene features on gene co-expression profiles

The potential effect of the three gene features, including mRNA abundance, gDNA size, and GC content, on gene co-expression profiles was first analyzed. The BrainSpan human brain transcriptome dataset was used for this analysis. This dataset contains transcriptomes of human (both sex) brain tissues from 16 different brain regions of various developmental stages and ages (from PCW8 to 40Y). A total of 12,250 genes with information on all 3 gene features were used for analyses. These genes were placed in ascending order of mRNA abundance, gDNA size, or GC content as ranked gene lists (**Table S1**). The correlation coefficient (CC) of each gene pair was calculated to reflect the co-expression level of the two genes, and results were displayed in pseudo-color-coded matrices. In each of the CC matrices (**Fig. 2A**), these 12,250 genes were placed in ascending order on both x and y axes. All three CC matrices exhibited variable color intensity in different areas with higher intensity corresponding to higher CC values. The overall color intensity was the highest in areas corresponding to medium mRNA abundance, medium to large gDNA size, and low GC content (**Fig. 2A**). This result suggests that all three gene features affect gene co-expression profiles.

**Figure 2.**
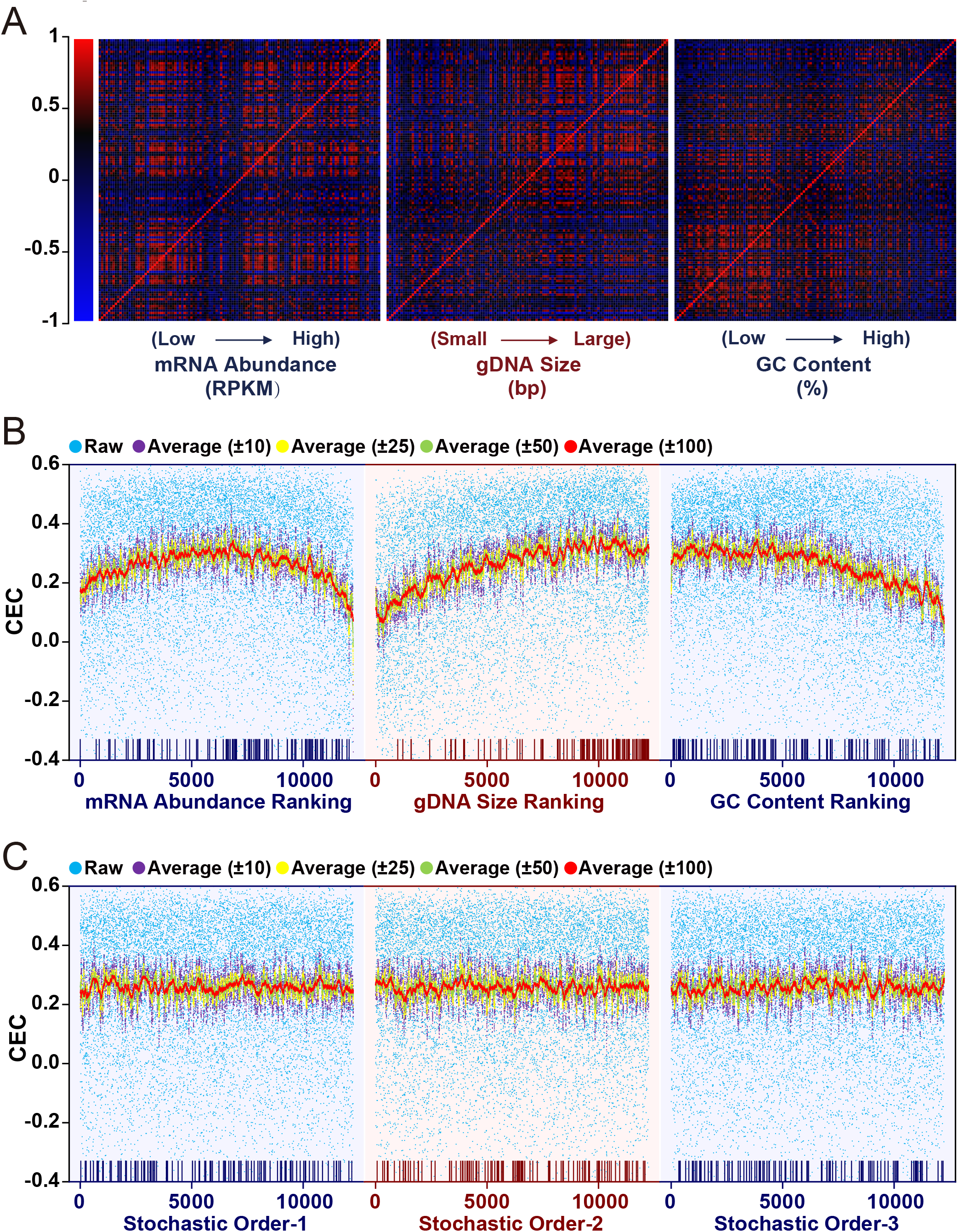
Effect of gene features on genome-wide gene co-expression profiles. A total of 12,250 genes with information on all 3 gene features were identified, placed in **Table S1** as gene lists, and used for analyses. **A**: Heatmaps of correlation coefficients (CCs) of genome-wide gene pairs. Genes were ranked according to mRNA abundance, gDNA size, or GC content. The correlation coefficient (CC) of each gene with all genes was plotted and displayed in pseudo-color-coded matrices. **B**: Genome-wide distribution of co-expression coefficients (CECs) of each gene with the hcASD gene set under three different gene ranking conditions. In each of the 3 matrix panels, the 12,250 genes were placed in ascending order on the x-axis, with 1 being the lowest mRNA abundance and GC content or shortest in gDNA size and 12,250 being the highest in mRNA abundance and GC content or the longest in gDNA size. Each blue dot represents the CEC of a gene with the hcASD gene set. Purple, yellow, green, and red dots represent noise-reduced (average) CEC of a gene with its neighboring 20 (10 above and 10 below; ± 10), 50 (± 25), 100 (± 50), or 200 (± 100) genes on the gene lists, respectively. Rods at the bottom of each panel show locations of hcASD genes on the ranked gene lists. **C**: Genome-wide distribution of CECs of each gene with the hcASD gene set when genes are placed in stochastic (random) orders.

Most hcASDs are large genes with medium to large mRNA abundance but with no apparent bias in GC content (**Fig. S1**). To determine whether each of these three gene features affects the co-expression of a gene with the hcASD gene set as a whole, the co-expression coefficient (CEC, mean CC between a gene and each of the hcASD genes) of each of the 12,250 genes with the entire hcASD gene set was calculated (blue dots in **Fig. 2B; Table S1**). In each of the 3 panels (**Fig. 2B)**, the 12,250 genes were placed in ascending order (x-axis). A noise-reduced (by data averaging) CEC distribution curve was then generated by plotting the average CEC of a gene with its neighboring 20 (10 above and 10 below; ±10), 50 (±25), 100 (±50), or 200 (±100) genes on the gene lists under each gene ranking condition. Results showed a bell-shaped curve when genes were ranked by mRNA abundance, suggesting that genes with medium expression levels are more likely to co-express with the hcASD gene set (**Fig. 2B**, left panel). There was an overall positive correlation between the gDNA size of a gene and its CEC with the hcASD gene set (**Fig. 2B**, middle panel). The CEC maintained a relatively high level (0.28 - 0.35) when the GC content ranged from low to medium level (approximately < 45%, x-axis < 6000) and then gradually declined with increasing GC content (**Fig. 2B**, right panel; **Table S1**).

With cubic regression, each noise-reduced CEC distribution curve was found to have an R^2^ value > 0.88 (**Fig. S2; Table S2**), indicating a significant correlation between each of these gene features and the tendency of co-expression of a gene with the hcASD gene set. When the 12,250 genes were placed in stochastic (random) orders, CECs were evenly distributed, and the noise-reduced CEC distribution curves were largely flat (**Fig. 2C**).

Similar genome-wide gene co-expression profiles of the hcASD gene set were observed in transcriptomes of early (8PCW - 2Y) and late (4Y - 40Y) stages (**Fig. S3A**), both sex, and different brain regions (**Fig. S3B, C**). These findings suggest that the co-expression profile of hcASD genes is affected by all three gene features, regardless of developmental stages, sex, and brain areas. Similar effects of these three gene features on the co-expression profiles of hcASD genes were observed when a set of 64 high susceptibility genes ^[24]^ were used as the hcASD gene set (**Fig. S4**).

### Similar co-expression profiles of feature-matched gene sets

The genome-wide gene co-expression profile of the hcASD gene set was then compared to the profiles of 200 feature-matched non-hcASD gene sets. Each gene set comprised an equal number (101) of randomly selected and feature-matched non-hcASD genes under the three different gene ranking conditions (**Fig. 3A**). These gene sets were named “matched random” (mRand) gene sets (see methods). In general, the genome-wide CEC distribution of hcASDs was similar to that of each of the 200 mRand gene sets under all three gene ranking conditions. These findings suggest that gene sets with matched gene features have a similar genome-wide co-expression profile as the hcASD gene set. However, genes with low to moderate mRNA abundance (approximately 1.2 - 30 RPMK, 1 - 10500 on x-axis) had higher noise-reduced CECs with the hcASD gene set than with any of the 200 mRand gene sets. In contrast, high mRNA abundance genes (> 30 RPKM; 10500 - 12250 on the x-axis) had lower noise-reduced CECs with the hcASD gene set than with most mRand gene sets. Moreover, genes with medium to large sizes had higher noise-reduced CECs with the hcASD gene set than with most size-matched mRand gene sets. Except those with the highest GC content, most genes had higher noise-reduced CECs with the hcASD gene set than with most GC content-matched mRand gene sets.

**Figure 3.**
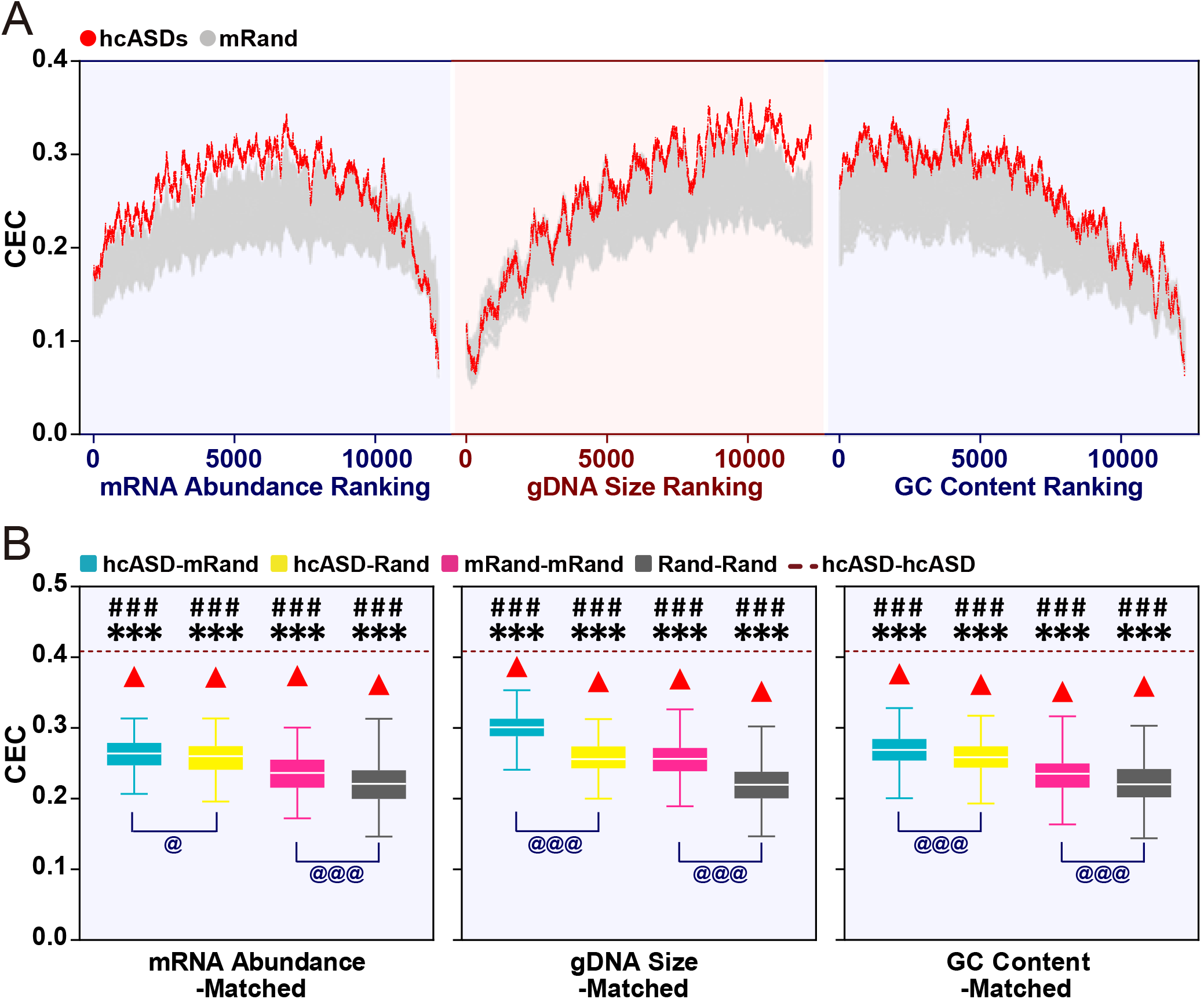
Convergent expression of hcASD genes determined by MGCA. **A**: Comparison of noise-reduced CEC distribution curves between the hcASD gene set and 200 matched random gene sets (mRand) under different gene ranking conditions. The X-axis represents gene ranks. **B**: CECs of hcASD-hcASD, hcASD-mRand, mRand-mRand, hcASD-Rand, and Rand-Rand gene set pairs. 200 each of mRand and Rand gene sets were analyzed. Box plots show ranges of CECs of hcASD-mRand, mRand-mRand, hcASD-Rand, and Rand-Rand gene set pairs. In each box plot, the central rectangles span the first quartile to the third quartile of 200 ranked CEC values. The white bar inside the rectangle shows the median CEC value, and whiskers above and below the box show the maximum and minimum values, respectively. The dotted line represents the CEC among hcASDs (hcASD-hcASD) in each panel. Three statistical methods were used to determine whether the CEC of hcASD-hcASD is significantly higher than that of hcASD-mRand, mRand-mRand, hcASD-Rand, and Rand-Rand. Upper fences test: red triangles stand for the boundaries of significant difference (3 x fence). Grubbs’ test: \*\*\**P* < 0.001. Permutation test: ### *P* < 0.001. The student’s *t*-test was used to determine whether the CECs of hcASD-mRand and mRand-mRand are significantly greater than those of hcASD-Rand and Rand-Rand, respectively. @ *P* < 0.05; @@@ *P* < 0.001.

### Co-expression of ASD risk genes

To determine whether hcASDs exhibit a significant tendency of co-expression with each other, the mean CEC of each of the 101 hcASDs with the hcASD gene set as a whole (hcASD-hcASD, see method) was compared to that of a large number of permuted gene sets, each comprised equal number of feature-matched non-hcASD genes (mRand-mRand) or randomly selected non-hcASD genes (Rand-Rand), and to the CEC between hcASD and mRand (hcASD-mRand) or Rand (hcASD-Rand) gene sets. Two hundred each of mRand and Rand gene sets were first analyzed. Results showed that feature-matched gene sets (mRand) had overall higher CECs than random gene sets (Rand) under all three matched conditions (@@@ in **Fig. 3B**), suggesting that genes with similar features tend to co-express with each other. The CEC of hcASD-hcASD (dashed line in **Fig. 3B**) was beyond the 3 times interquartile range [Q3 + 3 × (Q3 - Q1), 3x upper fence] of the CECs of mRand-mRand, Rand-Rand, hcASD-mRand, and hcASD-Rand gene sets. Results of Grubbs’ test confirmed this tendency (*** in **Fig. 3B**). These results suggest that hcASDs have a significantly greater co-expression tendency with each other than with other feature-matched non-hcASD genes or randomly selected genes. To corroborate this finding, a permutation test was conducted with 100,000 permuted sets of genes with matched or non-matched features. The CEC of hcASD-hcASD was again found to be significantly larger (permutation *P*-value < 0.00001) than that of hcASD-mRand, mRand-mRand, hcASD-Rand, or Rand-Rand (### in **Fig. 3B**), indicating a significant co-expression tendency of hcASDs.

Significant co-expression of hcASDs was also observed in transcriptomes of brain tissues from both early (8PCW – 2Y) and late (4Y – 40Y) stages (**Fig. S5A, B**), both sex (**Fig. S5C, D**), and different brain regions (**Fig. S6A-C**). These results indicate a highly conserved co-expression profile of hcASDs. Combined ranking of -log10 *P*-values of Grubbs’ test under all three different matched conditions was then performed to determine the relative significance level of co-expression of hcASDs with each other in different brain regions (**Fig. S6D**). The top four brain regions with the highest significance levels were cerebellum (CB), striatum (STR), orbital frontal cortex (OFC), and dorsal frontal cortex (DFC); these are the brain regions previously implicated in ASD ^[25–32]^. These results suggest that hcASDs play important roles in the development and function of these ASD-relevant brain regions.

### ASD-associated genes and pathways identified by MGCA

Genes whose CECs with hcASDs were significantly higher than with permuted gene sets composed of feature-matched genes under each of the three matched conditions were considered significantly co-expressed with hcASDs (estimated FDR of each gene below 1.25 x 10^-4^). These genes were named TriM (triple-matched) genes (**Figs. 1, 4A; Table S3**). TriM genes were then compared with a gene set containing 514 non-redundant genes from nine different sets of previously reported ASD-associated genes (cASD, see methods) and with a set of “True negative” ASD-associated genes, which were genes associated with non-mental health diseases but not with ASD ^[12]^. When the permutation *P*-value was either below 0.0001 or below 0.00001, the TriM gene set showed a significant enrichment of cASD genes (*P* = 0.0027 and P < 0.0001, respectively; chi-square test, **Fig. 4B**). This result suggests that at a permutation *P*-value below 0.0001, TriM genes have a high rate of positive prediction of being ASD-associated genes. When the permutation *P*-value was below 0.00001, TriM genes exhibited a significant negative enrichment of “True negative” genes (*P* = 0.0068; chi-square test, **Fig. 4B**), suggesting a significantly low false-positive rate in the prediction of ASD-associated genes.

**Figure 4.**
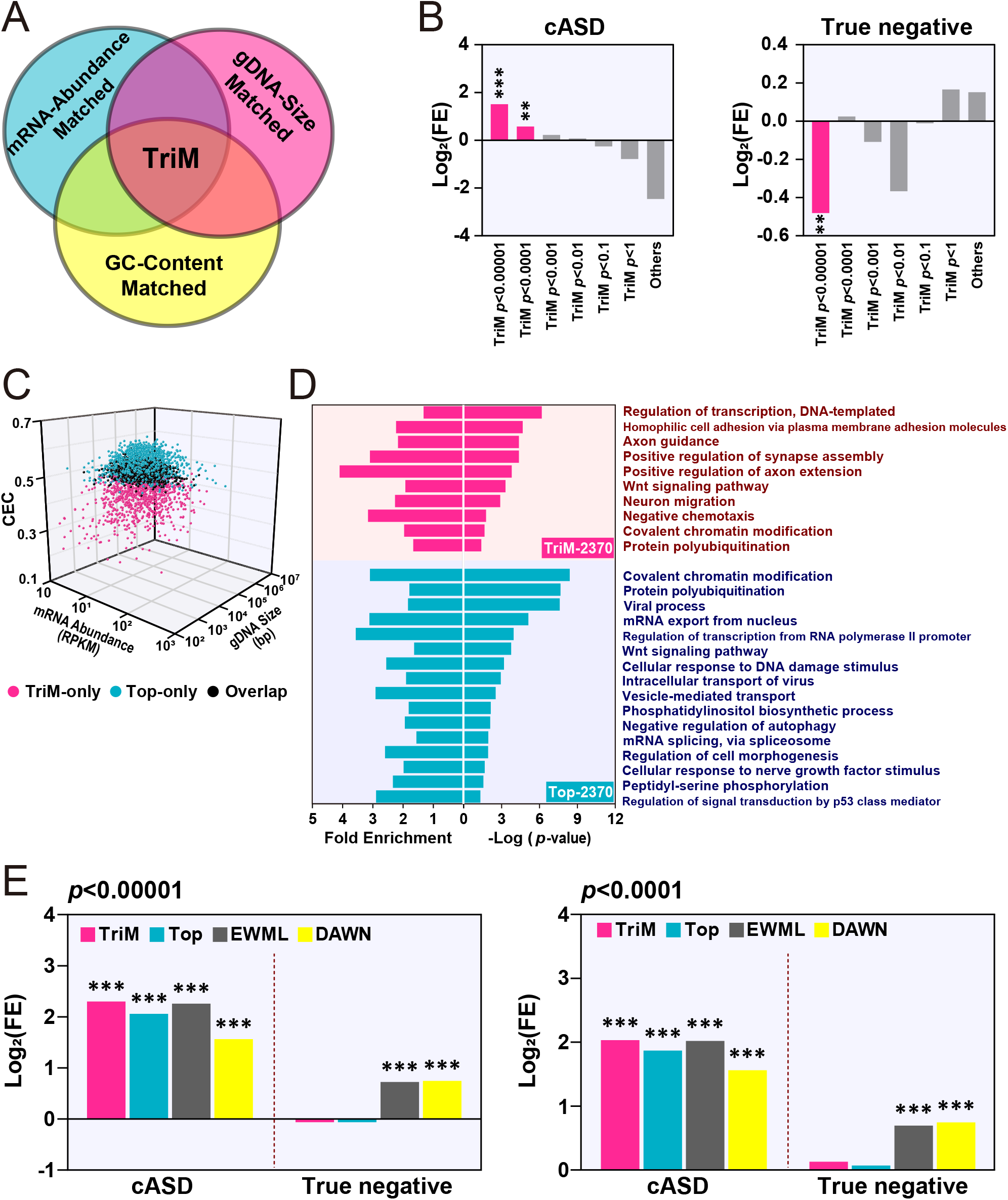
Evaluation of TriM genes identified by MGCA. **A**: Schematic of the TriM gene identification. TriM genes were significantly co-expressed with hcASDs under all three matched conditions determined by MGCA. **B**: Enrichment of ASD-associated genes and negative genes in TriM gene sets at different ranges of *P*-values. Fold enrichment (log_2_) of cASD (left) and “True negative” (right) genes in different groups of TriM genes are shown. ** *P* < 0.01, *** *P* < 0.001, chi-square test. **C**: Three-dimensional distribution of TriM genes (*P* < 0.0001) and an equal number (2370) of genes with highest CEC with hcASDs (Top). The three axes are CEC value, gDNA size (bp), and mRNA abundance (RPKM). Each dot in the graph represents a gene. The TriM-specific genes, Top-specific genes, and overlapped genes are shown in purple, blue, and black. **D**: Gene Ontology (GO) analysis of TriM (*P* < 0.0001) and Top (2370) genes. Significantly enriched GO terms are shown (*P* < 0.05, Benjamini-Hochberg correction). **E**: Comparison of the enrichment of cASD genes and “True negative” genes in TriM, Top, EWML-identified genes, and DAWN-identified genes. *** *P* < 0.001, chi-square test.

Altogether, MGCA revealed 2370 TriM genes with a permutation *P*-value < 0.0001. These genes (TriM-2370) were compared with an equal number (2370) of genes that had the highest CECs with the hcASD gene set (referred to as Top-2370 gene set, **Table S3**). TriM-2370 and Top-2370 gene sets had 1414 genes in common (overlapped), and each had 956 non-overlapped genes. These two non-overlapped gene sets were named TriM-only and Top-only, respectively (**Fig. 4C; Table S3**). Most Top-only genes had a medium mRNA abundance level, a large gene size, and a high CEC value (> 0.46), whereas TriM-only genes had a broad range of mRNA abundance, gene size, and CEC values (**Fig. 4C**). Gene ontology (GO) enrichment analysis of the TriM-2370 gene set showed significant over-representation of genes in molecular pathways closely related to ASD, including gene transcription regulation, homophilic cell adhesion, axon guidance and axon extension, synapse assembly, Wnt signaling pathway, neuron migration, covalent chromatin modification, and protein polyubiquitination. Fewer pathways relevant to ASD were revealed in the Top-2370 gene set by GO analysis, including covalent chromatin modification, transcription regulation, protein polyubiquitination, Wnt signaling pathway, and negative regulation of autophagy (**Fig. 4D; Table S4**). To investigate the functional relationship between cASD and TriM-2370 or Top-2370 gene sets, the enriched molecular pathways of cASD, TriM-only, Top-only, and overlapped gene sets were subject to pathway enrichment analysis ^[33]^ (**Fig. S7**). Results showed that TriM-only and Top-only gene sets converged in different but complementary molecular pathways. The molecular pathways of the cASD gene set were found to cluster closer to those of the TriM-only set than to those of the Top-only set, suggesting that TriM-only genes have a closer functional relationship with cASD genes than with Top-only genes. Some well-established ASD risk genes, such as *FOXP1*, *TBR1*, *SHANK2*, *SYNGAP1*, and *PCDH9*, were found in the TriM-only gene set, suggesting a better performance of MGCA than conventional GCA in revealing ASD-relevant molecular pathways.

The effectiveness of MGCA in predicting ASD-relevant genes was compared with that of GCA and two other risk gene prediction algorithms. One was DAWN (Detecting Association With Networks), which has been used to analyze the association between rare genetic variations and gene co-expression in the mid-fetal prefrontal and somatosensory cortex ^[34]^. The other algorithm was EWML (Evidence-Weighted Machine Learning) that has been used to predict the probability of ASD association with whole-genome genes based on data of gene co-expression, genetic mutations, and protein-protein interaction ^[12]^. The combined ASD risk gene set (cASD) and the “true negative” gene set were used to conduct the cross-comparison between different algorisms. At a permutation *P*-value of 0.0001 or 0.00001, TriM genes had higher enrichment of cASD genes than an equal number of genes with highest CEC values (Top), an equal number of top-ranked ASD-linked genes predicted by EWML, and network ASD genes (nASD) identified by DAWN (**Fig. 4E**). Furthermore, fewer TriM genes overlapped with “True negative” genes than genes predicted by DAWN and EWML, which had significant enrichment of “true negative” genes (**Fig. 4E**), suggesting a lower rate of false-positive prediction by MGCA than that by EWML and DAWN (**Fig. 4E**). Thus, MGCA performs better than GCA by a higher positive prediction rate and performs better than the EWML and DAWN algorisms by both a higher positive prediction rate and a lower prediction error.

### Co-expression of cadherin genes with hcASDs

Consistent with previous findings ^[8]^, we found that homophilic cell adhesion is the most significantly over-represented pathway of TriM-2370 genes (**Fig. 4D; Table S4**). Some cadherin family members in the TriM-2370 gene set, such as *PCDH19*, are known to be high-risk ASD genes (**Table S6**) that play important roles in brain circuit development ^[35, 36]^. Several cadherin family members were also found in the TriM-2370 gene set, including many members of the protocadherins β gene cluster and Dachsous Cadherin-related 1 (*DCHS1*), suggesting that these genes also participate in the development and function of ASD-relevant brain circuits. Some cadherin genes were not significantly co-expressed with hcASDs under any of the matched conditions; these genes were referred to as tri-negative genes (TriN; **Table S6**). Several recent genetic studies have implicated two type II cadherins, *CDH11* and *CDH9*, in ASD and other psychiatric diseases ^[37–41]^. The CEC values of *CDH11* and *CDH9* with hcASDs were ranked at 5244 and 9581, respectively, in whole-genome genes. Therefore, neither of them belonged top-ranked genes based on traditional GCA. Using MGCA, we found that *CDH11* and *CDH9* belonged to TriM and TriN gene sets, respectively. We thus hypothesized that *CDH11*, but not *CDH9*, is more likely to be associated with ASD.

### Autism-like traits of *Cdh11*-null mice

To assess the functional relevance of *CDH11* and *CDH9* to ASD, the behaviors of *Cdh11-* and *Cdh9*-null mice were investigated. In the open field test (OFT), both male and female *Cdh11*-null mice spent a longer time exploring the central area of the open field arena than wild type (WT) littermates (**Fig. 5A, D**). Heterozygous littermates showed a similar but less significant pattern. Total locomotion distance and average moving speed of *Cdh11*-null mice were slightly reduced compared to WT littermates (**Fig. 5B, C**). Both male and female *Cdh9*-null mice were largely normal in the OFT (**Fig. 5E-G**).

**Figure 5.**
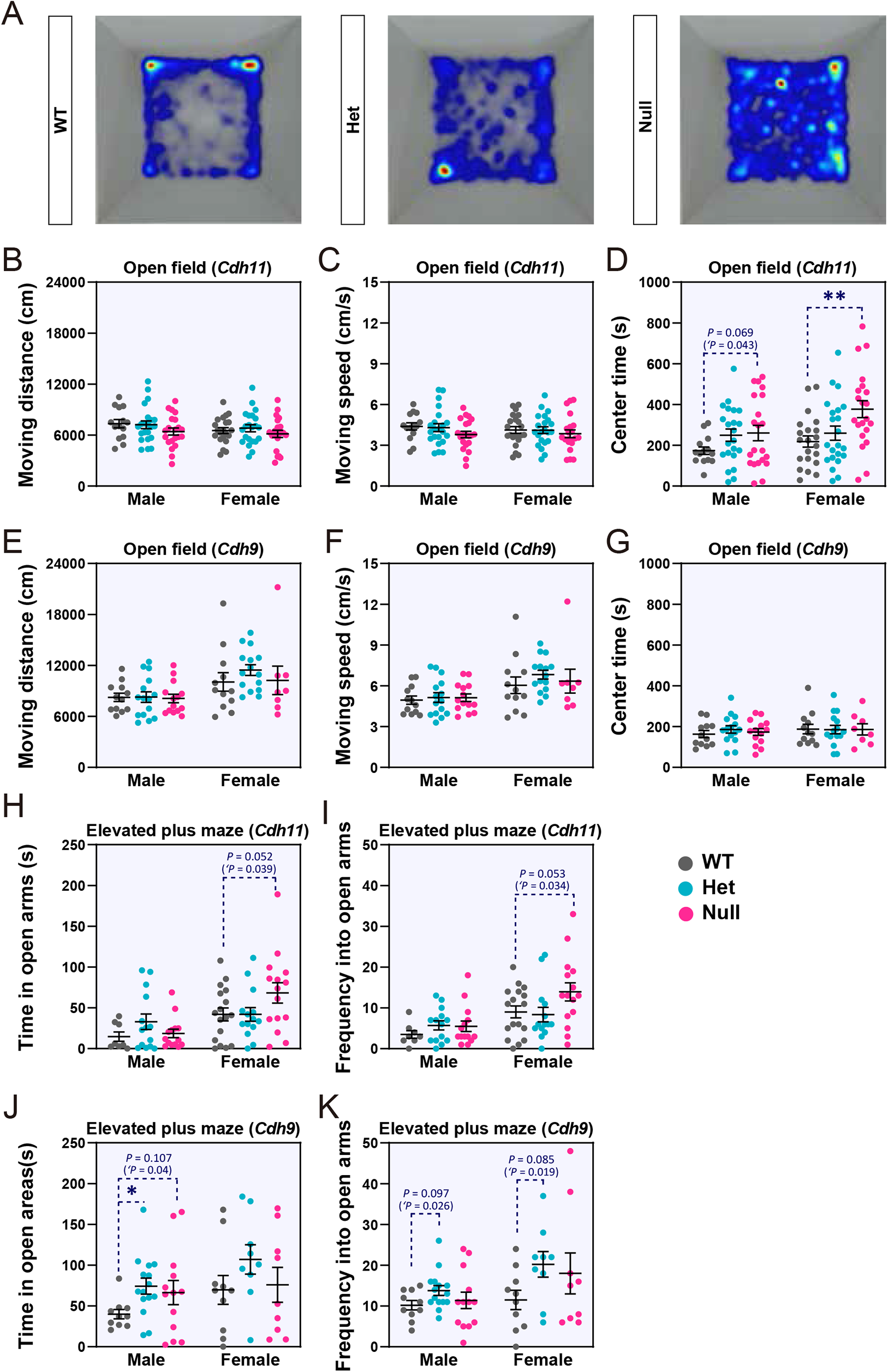
Open field and elevated plus maze tests of *Cdh11-* and *Cdh9*-null mice. **A**: Heatmaps showing cumulated frequency of locations visited by *Cdh11*-null mice, heterozygote (Het), and WT mice in the open field arena. **B, C,** and **D**: Moving distance, moving speed, and center exploration time of *Cdh11*-null mice. **E, F,** and **G**: Moving distance, moving speed, and center exploration time of *Cdh9*-null mice. (male *Cdh11*-null: n=21, Het: n=22, WT: n=14; female *Cdh11-*null: n=21, Het: n=22, WT: n=21; male *Cdh9*-null: n=14, Het: n=15, WT: n=12; female *Cdh9*-null: n=8, Het: n=15, WT: n=12). **J** and **K**: Time spent in open arm and open arm entry frequency of *Cdh11*-null mice. **H** and **I**: Time spent in open arm and open arm entry frequency of *Cdh9*-null mice (male *Cdh11*-null: n=14, Het: n=14, WT: n=8; female *Cdh11*-null: n=15, Het n=14, WT n=17; male *Cdh9*-null: n=13, Het: n=15, WT: n=10; female *Cdh9*-null: n=9, Het: n=9, WT: n=10). Data are mean ± SEM. Statistical difference was determined by Student *t*-test (‘*P* < 0.05) and by one-way ANOVA followed by Dunnett’s *t-*test (\**P* < 0.05, \*\**P* < 0.01).

In the elevated plus maze test, female *Cdh11*-null mice visited the open arm more frequently and spent a significantly longer time there. Heterozygous females spent a slightly but not statistically significantly more time in the open arm (**Fig. 5H, I**). The increased time and frequency of open arm exploration by female *Cdh11*-null mice is consistent with the results of a previous study using the same mouse line of mixed sex ^[42]^. Male *Cdh9*-null mice showed longer exploration of the open arm, but female *Cdh9*-null mice did not, although female heterozygotes showed an increased frequency of open arm entry (**Fig. 5J, K**).

Individuals with ASD often have a weaker grip strength than age-matched controls ^[43]^. The gripping strength test and the horizontal bar test showed that both male and female *Cdh11*-null mice exhibited significantly shorter hanging duration than WT littermates (**Fig. 6A, B**), indicating reduced gripping strength or impaired motor coordination. The gripping strength of *Cdh9*-null mice was normal (**Fig. 6C**).

**Figure 6.**
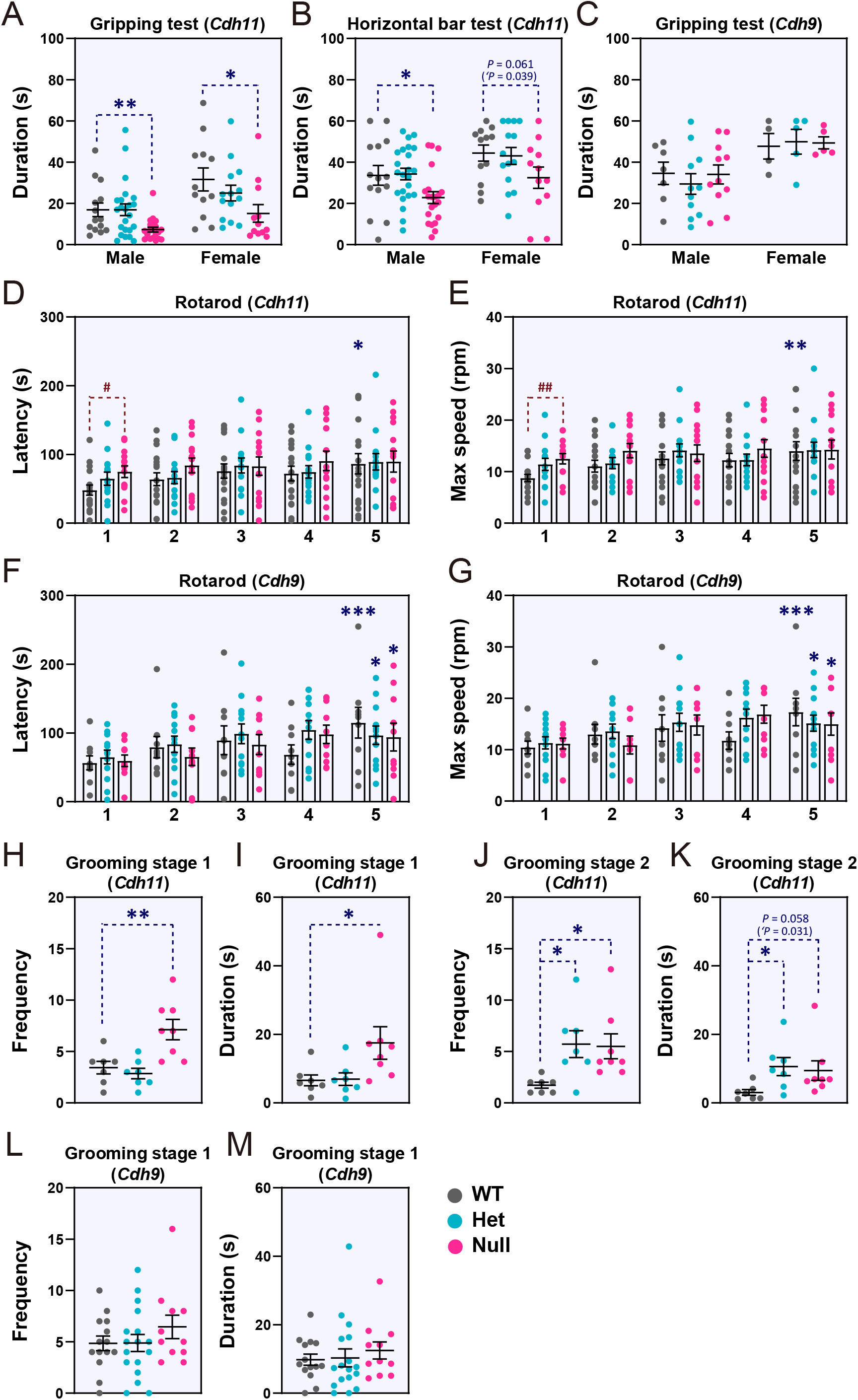
Gripping strength and repetitive behaviors of *Cdh11-* and *Cdh9*-null mice. **A** and **B**: Results of gripping test and horizontal bar test for *Cdh11*-null mice (male *Cdh11*-null: n=21, Het: n=23, WT: n=14; female *Cdh11*-null: n=12, Het: n=14, WT: n=12). **C**: Results of gripping test for *Cdh9*-null mice (male *Cdh9*-null: n=11, Het: n=11, WT: n=7, female *Cdh9*-null: n=5, Het: n=5, WT mice n=4). **D-G**: Latency to fall (**D, F**) and maximum durable speed (**E, G**) in rotarod test for female *Cdh11-* and *Cdh9*-null mice (*Cdh11*-null: n=14, Het: n=14, WT: n=17; *Cdh9*-null: n=10, Het: n=12; WT: n=9). Numbers below the X-axis (1-5) represent different trials of tests. **H-K**: Frequency and duration of self-grooming of female *Cdh11*-null mice during the first (stage 1, **H, I**) and the second (stage 2, **J, K**) 10 min in the open field arena (*Cdh11*-null: n=8, Het: n=7, WT n=7). **L** and **M,** Frequency and duration of self-grooming of female *Cdh9*-null mice during the first (stage 1) and second (stage 2) 10 min in the open filed arena (*Cdh9*-null: n=11, Het: n=17, WT: n=14). Data are mean ± SEM. Statistical difference was determined by Student *t*-test (‘*P* < 0.05) and by one-way ANOVA followed by Dunnett’s *t-*test (\**P* < 0.05, \*\**P* < 0.01, \*\*\**P* < 0.001, compared to first trial. # *P* < 0.05, ## *P* < 0.01, compared to mice of the WT littermates).

The rotarod test was conducted to evaluate motor-related functions of null mice. Since female and male mutant mice displayed similar behaviors in most of the above behavioral tests, only female mice were analyzed in this test. Compared to WT littermates, *Cdh11*-null mice, but not *Cdh9*-null mice, stayed longer on the rotarod and endured a higher rotation speed in the initial trial (**Fig. 6D-G**). In subsequent trials, *Cdh11*-null mice did not display significant performance improvement (**Fig. 6D, E**), indicating impaired motor learning. The enhanced performance of *Cdh11*-null mice in the initial trial was very similar to the phenotype of several other well-characterized ASD mouse models and suggested increased repetitive motion of these mutant mice ^[44]^.

Repetitive behaviors were then evaluated by measuring the duration and frequency of self-grooming within 10 minutes, during which mice were placed in a novel or a relatively familiar environment. As shown in **Fig. 6H-I**, during the first 10 minutes of exploring a novel chamber, *Cdh11*-null mice exhibited a significantly greater frequency of self-grooming than WT littermates, indicating elevated repetitive behavior in a novel environment. *Cdh11*-null mice also showed a significantly higher frequency of self-grooming than WT littermates during the second 10-minute period (**Fig. 6J, K**), indicating elevated repetitive behavior even in a relatively familiar environment. No such behavioral alteration was observed in *Cdh9*-null mice (**Fig. 6L, M**).

The modified three-chamber social preference test was conducted to evaluate the sociability of mutant mice. One main modification was an enlargement of the area for housing social partner mice to reduce their potential stress and anxiety. Another major modification to the protocol was using three mice instead of a single mouse as social partners. This was done to increase the availability of social cues and reduce the variability of test results caused by differences in the sociability of individual social partners (**Fig. 7A**). In addition, the two side chambers were covered on the top to slow the diffusion and mixing of odorant cues. Results showed that female *Cdh11*-null mice exhibited a significant preference to social partner mice than to an object and a significant preference to novel partners than to familiar ones (**Fig. 7B, C**). However, compared to WT littermates, mutant mice spent a significantly longer time in the middle chamber but significantly shorter time interacting with partner mice (**Fig. 7B, C**), indicating reduced sociability. In contrast, *Cdh9*-null mice did not show any abnormality in this test (**Fig. 7D, E**).

**Figure 7.**
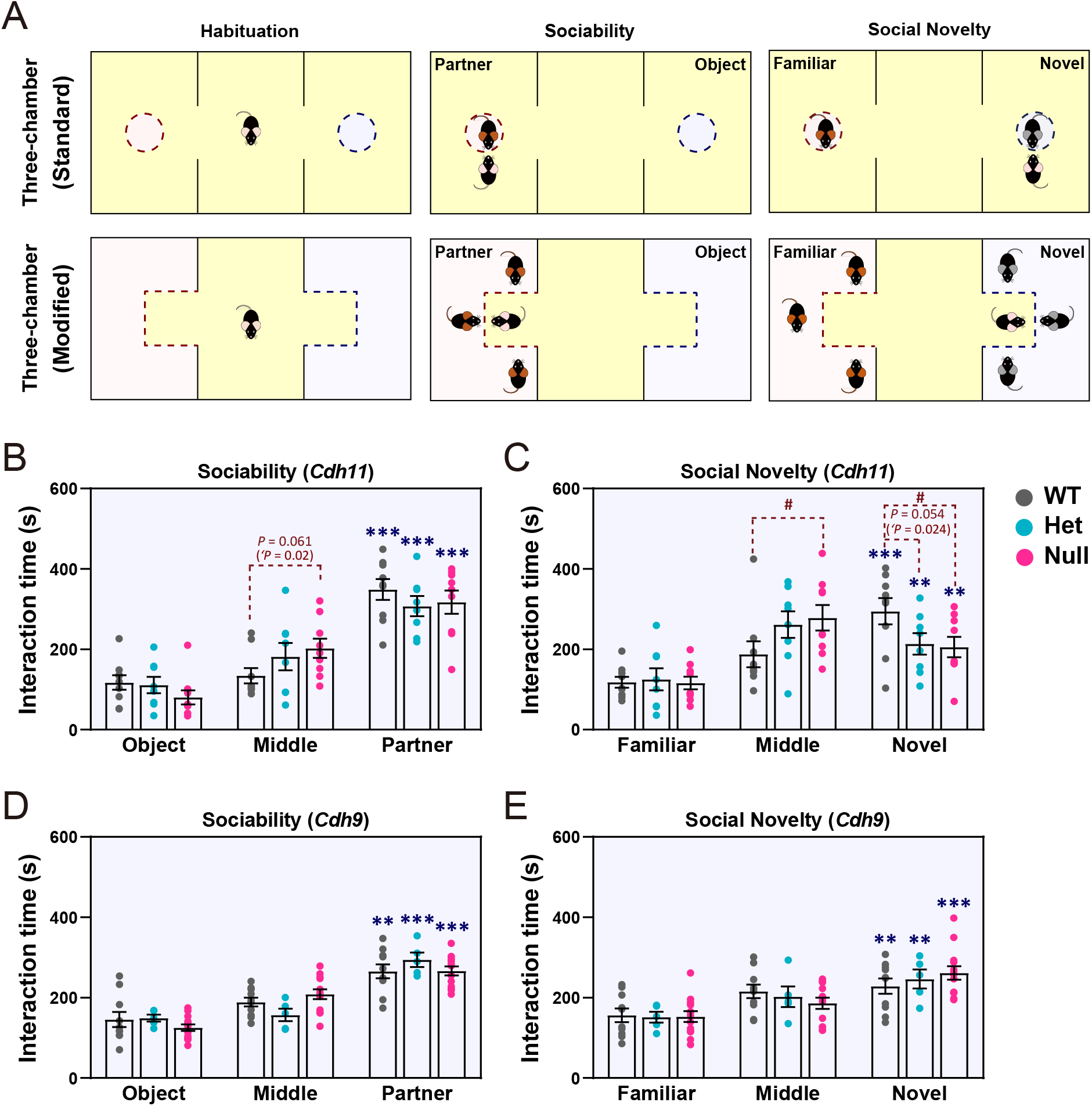
Modified three-chamber test of female *Cdh11-* and *Cdh9*-null mice. **a**: Schematics of standard and modified three-chamber tests. **b** and **c**: Results of sociability and social novelty preference tests of *Cdh11*-null mice (*Cdh11*-null: n=9, Het: n=8; WT: n=9). **d** and **e**: Results of sociability and novelty preference tests of *Cdh9*-null mice (*Cdh9*-null: n=13, Het: n=5, WT: n=10). Data are Mean ± SEM. Statistical difference was determined by Student *t*-test (‘*P* < 0.05) and by one-way ANOVA followed by Dunnett’s *t-*test (\**P* < 0.05, \*\**P* < 0.01, \*\*\**P* < 0.001, compared to the duration spent in the other side chamber. #*P* < 0.05, compared to WT littermates).

## Discussion

Gene co-expression analysis (GCA) is a powerful tool to find functionally convergent genes. Several previous GCA studies had considered the potential effect of gene size and GC content on the co-expression of ASD risk genes ^[7]^. In the present study, we discovered that three gene features, including mRNA abundance, gDNA size, and GC content, affected the genome-wide gene co-expression profiles in the brain. Although the mechanisms by which different gene features affect gene co-expression are unknown, our findings suggest the importance of considering the effect of these gene features in gene co-expression analysis (GCA). As an example of the potential influence of confounding gene features on GCA, genes that are stably expressed in the brain may have high CCs with each other. These aleatory high CCs of high-abundance genes do not mean a real co-expression relationship of genes of common molecular pathways. One possible influence of gene size on GCA is that large-size genes may have long noncoding regions that could be potential regulatory elements. Thus, compared to small genes, large genes may have more shared regulatory elements, which means a higher chance of being co-regulated by common transcription factors. Considering that most hcASD genes are large-size genes, they may have a higher tendency of co-expression with large genes. Genes with high GC content have higher mRNA stability and thus have higher chances of co-existence with each other. However, since most hcASD genes have low to medium GC content (**Fig 2B**), it may explain an overall negative correlation between the GC content of genes and their co-expression with hcASDs.

Almost all previous studies ignored these important confounding factors and just selected high-CC gene pairs to construct the gene co-expression network. Without correcting the effect of this confounding factor, the effectiveness of GCA would be compromised. Instead of setting a CC threshold for GCA as in most other studies, we screened for significant co-expression relationships by comparing the co-expression coefficient (CEC) of a gene with the hcASD gene set to that with permuted gene sets of matched gene features. Only genes that had a CEC with the hcASD gene set significantly higher than its CECs with permuted sets of feature-matched genes were considered being co-expressed with hcASDs. This matched-gene co-expression analysis (MGCA) paradigm **(Fig. 1)** allowed the demonstration of significant co-expression of hcASDs and avoided the potential bias caused by an empirically determined threshold for CC of gene pairs in GCA. Our results revealed that MGCA is more efficient in predicting gene association than the pre-existing methods DAWN, an algorism integrating genetic variants and gene co-expression data, and EWML, a sophisticated machine learning algorithm with the integration of gene co-expression, gene mutation database, and PPI network. We believe that the high performance of MGCA could be attributed to the correction of three confounding gene features in the determination of functionally relevant gene co-expression. Although correlations of these confounding gene features with the co-expression with hcASDs have been considered ^[7]^, the potential interference of these gene features on the construction of gene co-expression networks was not considered in previous studies. Therefore, MGCA will be an important complement to current gene association prediction algorithms (**Fig. 1**). As MGCA is based solely on the gene co-expression data, future algorithms combining MGCA with genetic mutation data and machine learning will further improve its efficacy.

An important finding in this study is the plausible association of *CDH11* with ASD determined by MGCA. Cadherins have been shown to accumulate in synaptic junctions and regulate dendrite development and synapse maturation ^[45–48]^. Several cadherin family members, such as some protocadherins in the *FAT* cadherin subfamily, have been implicated in ASD ^[49–58]^. A genetic association study of a large cohort of ASD individuals and matched controls revealed genes in the protocadherin α gene cluster (*PCDHA*) as ASD risk genes^[51]^. Mutations in the *PCDH19* gene have been shown to cause early-onset epilepsy, and many individuals with these mutations also display autistic features ^[52–54]^. Mutations in the cadherin EGF LAG seven-pass G-type receptor 2 gene (*CELSR2*) were speculated to be responsible for the Joubert syndrome, a disease with a high degree of autistic features ^[59, 60]^. It is uncertain whether other cadherins are also high-risk factors. Using MGCA, we found that a group of cadherin superfamily members exhibited a high co-expression with hcASDs, suggesting shared functions with hcASDs and a role in ASD etiology. Among them, several protocadherins, mainly *PCDHB*s, exhibited significant co-expression with hcASDs (**Table S6**). The functions of these putative ASD-associated cadherins in the brain remain to be determined. One of such cadherins identified by MGCA is CDH11. In this study, we found that *Cdh11*-null mice had significantly increased repetitive behaviors. The brain regions, including the neocortex, CB, and STR, are known to be involved in the control of repetitive behaviors ^[61]^. It is likely that cadherins, *Cdh11* in particular, play important roles in mediating synapse formation during the wiring of circuits in these brain areas. Consistent with this postulation, our recent work showed *Cdh11* expression in ASD-associated sub-regions in the CB of developing mouse brain ^[62]^.

In human studies, partial deletion of *CDH11* was observed in a sporadic case of non-syndromic ASD, mild intellectual disability, and attention deficit hyperactivity disorder (ADHD) ^[37]^. A case-control association study revealed a high prevalence of a homozygous single nucleotide variant rs7187376C/C of *CDH11* in patients with ASD ^[37]^. Several other coding variants of *CDH11* were also discovered in ASD individuals ^[37]^. Behavioral changes that we have observed in *Cdh11*-null mice, including reduced anxiety, increased repetitive behavior, and reduced sociability, are highly consistent with the non-syndromic ASD case with partial deletion of *CDH11*^[37]^. This observation supports the notion that loss-of-function of a single gene, such as *CDH11*, is sufficient to cause major autism traits. Recessive mutations have been implicated in ASD, and biallelic disruption of recessive neurodevelopmental genes in ASD has been reported ^[62]^. We found that homozygous *Cdh11*-null mice, but not heterozygous *Cdh11*-null mice, displayed autism-like behavioral deficits. This suggests that *CDH11* may be a recessive gene of ASD. However, clinical data from more families with *CDH11* mutations are needed to determine whether it is true in the future. Behavioral phenotypes of ASD are highly heterogeneous. Some individuals with ASD are hypoactive with elevated anxiety, and some have attention deficit hyperactivity disorder (ADHD) but with reduced anxiety ^[64–67]^. The genetic and neurobiological mechanisms underlying this behavioral heterogeneity have not been fully determined. Further investigation with a larger cohort of patient families is needed to determine whether loss-of-function mutations of *CDH11* are associated with ADHD.

Most genetic variants found in patients with ASD are heterozygous. In some behavioral tests, heterozygous *Cdh11*-null mice showed a similar trend of behavioral alterations as homozygous null mice, but not at a statistically significant level (**Figs. 6J, K; 7C**). As ASD has a complex genetic basis and is affected by environmental factors, it is conceivable that the haplodeficiency of a single risk gene causes a relatively mild behavioral phenotype in mice. More severe behavioral deficits may result if the haplodeficiency of *Cdh11* is combined with other genetic or environmental factors. Our findings suggest that *CDH11* is significantly co-expressed with hcASDs and that its mutations may exert a causal effect in autism traits. *Cdh11*-null mice would be very useful in dissecting the circuit mechanisms underlying a subgroup of ASD and in screening drugs targeting this subgroup of ASD.

*CDH9* plays a vital role in establishing specific synaptic wiring in both the hippocampus and the retina ^[22, 68]^. Its association with ASD has been suggested by several studies on exome sequencing ^[49, 69]^. The primary evidence linking *CDH9* to ASD is the strong association of the single nucleotide polymorphism rs4307059 located in the intergenic region between *CDH10* and *CDH9* with ASD ^[70]^. However, this rs4307059 genotype was not correlated with the expression of either *CDH9* or *CDH10* in adult brains ^[70, 71]^, and whether a correlation exists in fetal brains is unknown. Recently, an antisense noncoding RNA of the moesin pseudogene 1 (MSNP1AS) was shown to be transcribed from the locus harboring rs4307059. Alterations in this pseudogene were postulated to contribute to ASD ^[71–73]^. Whether *CDH9* deficiency is a causal factor for ASD remains undetermined. Our MGCA showed that, unlike *CDH11*, *CDH9* was not co-expressed with hcASDs. This is an indication that *CDH9* may not play an essential role in the wiring of ASD-relevant circuits. Consistent with this notion, behavioral tests showed that *Cdh9*-null mice exhibited a very mild behavioral abnormality only in the elevated plus maze test but not in any other tests. With recent findings by other researchers ^[74]^, our results suggest that *CDH9* deficiency may not have a significant effect on autism traits.

In conclusion, this study revealed the importance of considering matched gene features in the analysis of gene co-expression and demonstrated the effectiveness of MGCA in the identification of putative ASD-associated genes and their convergent signaling pathways. The application of MGCA led to the determination of *CDH11* as a putative ASD-associated gene. Our results also showed that *Cdh11*-null mice could be used to study circuit mechanisms of a subgroup of ASD and explore therapeutic strategies for ASD.

## Acknowledgments

We thank Elizabeth Benevides for editing the manuscript. This work was funded by the National Science Foundation of China (NSFC-31871501, NSFC-ISF-32061143016) and Hussman Foundation (HIAS15006). X.Y. was supported by the Simons Foundation (296143).

## Authors’ contributions

N.W., Y.W., and Y.P. analyzed data and generated figures. J.J and X.Y. conducted behavioral tests. Y.P. and X.Y. designed the study and wrote the manuscript.

## Supplementary Information

**Supplemental_Fig. S1**

Distribution of hcASDs in gene rank matrices. Genes in the whole genome were ranked by gDNA size, GC content (horizontal axis), and mRNA abundance level (vertical axis) and plotted in matrices. Each blue dot represents a single gene, and each red dot represents an hcASD gene. Values of both horizontal and vertical axes are ranks (orders) of the 12,250 genes under 3 different gene ranking conditions.

**Supplemental_Fig. S2**

Fitting (regression) of CEC distribution curves under three different gene ranking conditions.

**Supplemental_Fig. S3**

CEC distribution curves for different developmental stages, sex, and brain regions.

**Supplemental_Fig. S4**

Genome-wide distribution of CECs of each gene with the hcASD (64) gene set under three different gene ranking conditions.

**Supplemental_Fig. S5**

Co-expression of hcASDs in the brain of different developmental stages and sex. **A** and **B** show data from early (8PCW - 2Y) and late (3Y - 40Y) developmental stages, respectively. **c** and **d** show data from male and female brain tissues, respectively.

**Supplemental_Fig. S6**

Co-expression of hcASDs in different brain regions. **A-C:** Box plots show the range of CECs of hcASD-mRand in different brain regions. **D:** Significant scores of co-expression of hcASDs in different brain regions. Score = N - R. N: number of brain regions. R: integrated ranking (average of 3 conditions) of -Log(*P*-value) in Grubbs’ test under four different gene ranking conditions.

**Supplemental_Fig. S7**

Clustering analysis of –Log(*P*) values in GO analysis of TriM-only, Top-only, Overlapped, and combined ASD gene sets (cASDs). –Log(*P*) values (0 - 20) are color-coded and shown in the heatmap.

**Supplemental_Table_S1**

List of total genes and hcASDs included in this study.

**Supplemental_Table_S2**

Parameters of fitted curves (regression curves) in supplemental_Fig_S2.

**Supplemental_Table_S3**

*P*-value and FDR of each gene under different gene ranking conditions and lists of TriM, Top, TriM-only, Top-only, and Overlap genes.

**Supplemental_Table_S4**

GO entries of TriM (*P* < 0.0001) and Top (2370) genes.

**Supplemental_Table_S5**

Lists of cASDs, True negative genes, EWML genes, and DAWN genes.

**Supplemental_Table_S6**

List of TriM- and TriN-cadherins

## References

[1] Berg JM, Geschwind DH. Autism genetics: searching for specificity and convergence. Genome Biol 2012, 13: 247.

[2] Jeste SS, Geschwind DH. Disentangling the heterogeneity of autism spectrum disorder through genetic findings. Nature Reviews Neurology 2014, 10: 74–81.

[3] de la Torre-Ubieta L, Won H, Stein JL, Geschwind DH. Advancing the understanding of autism disease mechanisms through genetics. Nat Med 2016, 22: 345–361.

[4] Sullivan PF, Geschwind DH. Defining the Genetic, Genomic, Cellular, and Diagnostic Architectures of Psychiatric Disorders. Cell 2019, 177: 162–183.

[5] Ronemus M, Iossifov I, Levy D, Wigler M. The role of de novo mutations in the genetics of autism spectrum disorders. Nat Rev Genet 2014, 15: 133–141.

[6] Courchesne E, Pramparo T, Gazestani VH, Lombardo MV, Pierce K, Lewis NE. The ASD Living Biology: from cell proliferation to clinical phenotype. Mol Psychiatry 2019, 24: 88–107.

[7] Willsey AJ, Sanders SJ, Li M, Dong S, Tebbenkamp AT, Muhle RA, et al. Coexpression networks implicate human midfetal deep cortical projection neurons in the pathogenesis of autism. Cell 2013, 155: 997–1007.

[8] Parikshak NN, Luo R, Zhang A, Won H, Lowe JK, Chandran V, et al. Integrative functional genomic analyses implicate specific molecular pathways and circuits in autism. Cell 2013, 155: 1008–1021.

[9] Stuart JM, Segal E, Koller D, Kim SK. A gene-coexpression network for global discovery of conserved genetic modules. Science 2003, 302: 249–255.

[10] Hormozdiari F, Penn O, Borenstein E, Eichler EE. The discovery of integrated gene networks for autism and related disorders. Genome Res 2015, 25: 142–154.

[11] Mahfouz A, Ziats MN, Rennert OM, Lelieveldt BP, Reinders MJ. Shared Pathways Among Autism Candidate Genes Determined by Co-expression Network Analysis of the Developing Human Brain Transcriptome. J Mol Neurosci 2015, 57: 580–594.

[12] Krishnan A, Zhang R, Yao V, Theesfeld CL, Wong AK, Tadych A, et al. Genome-wide prediction and functional characterization of the genetic basis of autism spectrum disorder. Nat Neurosci 2016, 19: 1454–1462.

[13] Kopp N, Climer S, Dougherty JD. Moving from capstones toward cornerstones: successes and challenges in applying systems biology to identify mechanisms of autism spectrum disorders. Front Genet 2015, 6: 301.

[14] Gabel HW, Kinde B, Stroud H, Gilbert CS, Harmin DA, Kastan NR, et al. Disruption of DNA-methylation-dependent long gene repression in Rett syndrome. Nature 2015, 522: 89–93.

[15] Satterstrom FK, Kosmicki JA, Wang JB, Breen MS, De Rubeis S, An JY, et al. Large-Scale Exome Sequencing Study Implicates Both Developmental and Functional Changes in the Neurobiology of Autism. Cell 2020, 180: 568–584.

[16] Abrahams BS, Arking DE, Campbell DB, Mefford HC, Morrow EM, Weiss LA, et al. SFARI Gene 2.0: a community-driven knowledgebase for the autism spectrum disorders (ASDs). Mol Autism 2013, 4: 36.

[17] SCEa, Consortium S. SPARK: A US Cohort of 50,000 Families to Accelerate Autism Research. Neuron 2018, 97: 488–493.

[18] Basu SN, Kollu R, Banerjee-Basu S. AutDB: a gene reference resource for autism research. Nucleic Acids Res 2009, 37: D832–836.

[19] RK CY, Merico D, Bookman M, J LH, Thiruvahindrapuram B, Patel RV, et al. Whole genome sequencing resource identifies 18 new candidate genes for autism spectrum disorder. Nat Neurosci 2017, 20: 602–611.

[20] Iossifov I, O’Roak BJ, Sanders SJ, Ronemus M, Krumm N, Levy D, et al. The contribution of de novo coding mutations to autism spectrum disorder. Nature 2014, 515: 216–221.

[21] Lin Y, Afshar S, Rajadhyaksha AM, Potash JB, Han S. A Machine Learning Approach to Predicting Autism Risk Genes: Validation of Known Genes and Discovery of New Candidates. Front Genet 2020, 11: 500064.

[22] Duan X, Krishnaswamy A, De la Huerta I, Sanes JR. Type II cadherins guide assembly of a direction-selective retinal circuit. Cell 2014, 158: 793–807.

[23] Horikawa K, Radice G, Takeichi M, Chisaka O. Adhesive subdivisions intrinsic to the epithelial somites. Dev Biol 1999, 215: 182–189.

[24] Sanders SJ, Xin H, Willsey AJ, Ercan-Sencicek AG, Samocha KE, Cicek AE, et al. Insights into Autism Spectrum Disorder Genomic Architecture and Biology from 71 Risk Loci. Neuron 2015, 87: 1215–1233.

[25] Tsai PT, Hull C, Chu Y, Greene-Colozzi E, Sadowski AR, Leech JM, et al. Autistic-like behaviour and cerebellar dysfunction in Purkinje cell Tsc1 mutant mice. Nature 2012, 488: 647–651.

[26] Khan AJ, Nair A, Keown CL, Datko MC, Lincoln AJ, Muller RA. Cerebro-cerebellar Resting-State Functional Connectivity in Children and Adolescents with Autism Spectrum Disorder. Biol Psychiatry 2015, 78: 625–634.

[27] Buxhoeveden DP, Semendeferi K, Buckwalter J, Schenker N, Switzer R, Courchesne E. Reduced minicolumns in the frontal cortex of patients with autism. Neuropathol Appl Neurobiol 2006, 32: 483–491.

[28] Green SA, Hernandez L, Tottenham N, Krasileva K, Bookheimer SY, Dapretto M. Neurobiology of Sensory Overresponsivity in Youth With Autism Spectrum Disorders. JAMA Psychiatry 2015, 72: 778–786.

[29] Green SA, Rudie JD, Colich NL, Wood JJ, Shirinyan D, Hernandez L, et al. Overreactive brain responses to sensory stimuli in youth with autism spectrum disorders. J Am Acad Child Adolesc Psychiatry 2013, 52: 1158–1172.

[30] Di Martino A, Kelly C, Grzadzinski R, Zuo XN, Mennes M, Mairena MA, et al. Aberrant striatal functional connectivity in children with autism. Biol Psychiatry 2011, 69: 847–856.

[31] Wang X, Bey AL, Katz BM, Badea A, Kim N, David LK, et al. Altered mGluR5-Homer scaffolds and corticostriatal connectivity in a Shank3 complete knockout model of autism. Nat Commun 2016, 7: 11459.

[32] Li X, Zhang K, He X, Zhou J, Jin C, Shen L, et al. Structural, Functional, and Molecular Imaging of Autism Spectrum Disorder. Neurosci Bull 2021.

[33] Zhou Y, Zhou B, Pache L, Chang M, Khodabakhshi AH, Tanaseichuk O, et al. Metascape provides a biologist-oriented resource for the analysis of systems-level datasets. Nat Commun 2019, 10: 1523.

[34] Liu L, Lei J, Sanders SJ, Willsey AJ, Kou Y, Cicek AE, et al. DAWN: a framework to identify autism genes and subnetworks using gene expression and genetics. Molecular Autism 2014, 5(1): 22.

[35] Yin J, Chun CA, Zavadenko NN, Pechatnikova NL, Naumova OY, Doddapaneni HV, et al. Next Generation Sequencing of 134 Children with Autism Spectrum Disorder and Regression. Genes (Basel) 2020, 11(8): 853.

[36] Emond MR, Biswas S, Jontes JD. Protocadherin-19 is essential for early steps in brain morphogenesis. Developmental Biology 2009, 334: 72–83.

[37] Crepel A, De Wolf V, Brison N, Ceulemans B, Walleghem D, Peuteman G, et al. Association of CDH11 with non-syndromic ASD. Am J Med Genet B Neuropsychiatr Genet 2014, 165B: 391–398.

[38] Gasso P, Ortiz AE, Mas S, Morer A, Calvo A, Bargallo N, et al. Association between genetic variants related to glutamatergic, dopaminergic and neurodevelopment pathways and white matter microstructure in child and adolescent patients with obsessive-compulsive disorder. J Affect Disord 2015, 186: 284–292.

[39] Sokolowski M, Wasserman J, Wasserman D. Polygenic associations of neurodevelopmental genes in suicide attempt. Mol Psychiatry 2016, 21: 1381–1390.

[40] Wang K, Zhang H, Bloss CS, Duvvuri V, Kaye W, Schork NJ, et al. A genome-wide association study on common SNPs and rare CNVs in anorexia nervosa. Mol Psychiatry 2011, 16: 949–959.

[41] Anazi S, Maddirevula S, Faqeih E, Alsedairy H, Alzahrani F, Shamseldin HE, et al. Clinical genomics expands the morbid genome of intellectual disability and offers a high diagnostic yield. Mol Psychiatry 2017, 22: 615–624.

[42] Manabe T, Togashi H, Uchida N, Suzuki SC, Hayakawa Y, Yamamoto M, et al. Loss of cadherin-11 adhesion receptor enhances plastic changes in hippocampal synapses and modifies behavioral responses. Mol Cell Neurosci 2000, 15: 534–546.

[43] Travers BG, Bigler ED, Tromp do PM, Adluru N, Destiche D, Samsin D, et al. Brainstem White Matter Predicts Individual Differences in Manual Motor Difficulties and Symptom Severity in Autism. J Autism Dev Disord 2015, 45: 3030–3040.

[44] Rothwell PE, Fuccillo MV, Maxeiner S, Hayton SJ, Gokce O, Lim BK, et al. Autism-associated neuroligin-3 mutations commonly impair striatal circuits to boost repetitive behaviors. Cell 2014, 158: 198–212.

[45] Bekirov IH, Nagy V, Svoronos A, Huntley GW, Benson DL. Cadherin-8 and N-cadherin differentially regulate pre- and postsynaptic development of the hippocampal mossy fiber pathway. Hippocampus 2008, 18: 349–363.

[46] Friedman LG, Riemslagh FW, Sullivan JM, Mesias R, Williams FM, Huntley GW, et al. Cadherin-8 expression, synaptic localization, and molecular control of neuronal form in prefrontal corticostriatal circuits. J Comp Neurol 2015, 523: 75–92.

[47] Bian WJ, Miao WY, He SJ, Qiu Z, Yu X. Coordinated Spine Pruning and Maturation Mediated by Inter-Spine Competition for Cadherin/Catenin Complexes. Cell 2015, 162: 808–822.

[48] Tsai NP, Wilkerson JR, Guo W, Maksimova MA, DeMartino GN, Cowan CW, et al. Multiple autism-linked genes mediate synapse elimination via proteasomal degradation of a synaptic scaffold PSD-95. Cell 2012, 151: 1581–1594.

[49] Cukier HN, Dueker ND, Slifer SH, Lee JM, Whitehead PL, Lalanne E, et al. Exome sequencing of extended families with autism reveals genes shared across neurodevelopmental and neuropsychiatric disorders. Mol Autism 2014, 5: 1.

[50] Butler MG, Rafi SK, Hossain W, Stephan DA, Manzardo AM. Whole exome sequencing in females with autism implicates novel and candidate genes. Int J Mol Sci 2015, 16: 1312–1335.

[51] Anitha A, Thanseem I, Nakamura K, Yamada K, Iwayama Y, Toyota T, et al. Protocadherin alpha (PCDHA) as a novel susceptibility gene for autism. J Psychiatry Neurosci 2013, 38: 192–198.

[52] Cappelletti S, Specchio N, Moavero R, Terracciano A, Trivisano M, Pontrelli G, et al. Cognitive development in females with PCDH19 gene-related epilepsy. Epilepsy Behav 2015, 42: 36–40.

[53] van Harssel JJ, Weckhuysen S, van Kempen MJ, Hardies K, Verbeek NE, de Kovel CG, et al. Clinical and genetic aspects of PCDH19-related epilepsy syndromes and the possible role of PCDH19 mutations in males with autism spectrum disorders. Neurogenetics 2013, 14: 23–34.

[54] Camacho A, Simon R, Sanz R, Vinuela A, Martinez-Salio A, Mateos F. Cognitive and behavioral profile in females with epilepsy with PDCH19 mutation: two novel mutations and review of the literature. Epilepsy Behav 2012, 24: 134–137.

[55] Sanders SJ, Ercan-Sencicek AG, Hus V, Luo R, Murtha MT, Moreno-De-Luca D, et al. Multiple recurrent de novo CNVs, including duplications of the 7q11.23 Williams syndrome region, are strongly associated with autism. Neuron 2011, 70: 863–885.

[56] Pagnamenta AT, Khan H, Walker S, Gerrelli D, Wing K, Bonaglia MC, et al. Rare familial 16q21 microdeletions under a linkage peak implicate cadherin 8 (CDH8) in susceptibility to autism and learning disability. J Med Genet 2011, 48: 48–54.

[57] Moya PR, Dodman NH, Timpano KR, Rubenstein LM, Rana Z, Fried RL, et al. Rare missense neuronal cadherin gene (CDH2) variants in specific obsessive-compulsive disorder and Tourette disorder phenotypes. Eur J Hum Genet 2013, 21: 850–854.

[58] Redies C, Hertel N, Hubner CA. Cadherins and neuropsychiatric disorders. Brain Res 2012, 1470: 130–144.

[59] Holroyd S, Reiss AL, Bryan RN. Autistic features in Joubert syndrome: a genetic disorder with agenesis of the cerebellar vermis. Biol Psychiatry 1991, 29: 287–294.

[60] Feng J, Han Q, Zhou L. Planar cell polarity genes, Celsr1-3, in neural development. Neurosci Bull 2012, 28: 309–315.

[61] Kim H, Lim CS, Kaang BK. Neuronal mechanisms and circuits underlying repetitive behaviors in mouse models of autism spectrum disorder. Behav Brain Funct 2016, 12: 3.

[62] Wang C, Pan YH, Wang Y, Blatt G, Yuan XB. Segregated expressions of autism risk genes *Cdh11* and *Cdh9* in autism-relevant regions of developing cerebellum. Mol Brain 2019, 12: 40.

[63] Doan RN, Lim ET, De Rubeis S, Betancur C, Cutler DJ, Chiocchetti AG, et al. Recessive gene disruptions in autism spectrum disorder. Nat Genet 2019, 51: 1092–1098.

[64] Gadow KD, Devincent CJ, Pomeroy J, Azizian A. Comparison of DSM-IV symptoms in elementary school-age children with PDD versus clinic and community samples. Autism 2005, 9: 392–415.

[65] Simonoff E, Pickles A, Charman T, Chandler S, Loucas T, Baird G. Psychiatric disorders in children with autism spectrum disorders: prevalence, comorbidity, and associated factors in a population-derived sample. J Am Acad Child Adolesc Psychiatry 2008, 47: 921–929.

[66] Greene RW, Biederman J, Faraone SV, Ouellette CA, Penn C, Griffin SM. Toward a new psychometric definition of social disability in children with attention-deficit hyperactivity disorder. J Am Acad Child Adolesc Psychiatry 1996, 35: 571–578.

[67] Grzadzinski R, Dick C, Lord C, Bishop S. Parent-reported and clinician-observed autism spectrum disorder (ASD) symptoms in children with attention deficit/hyperactivity disorder (ADHD): implications for practice under DSM-5. Mol Autism 2016, 7: 7.

[68] Williams ME, Wilke SA, Daggett A, Davis E, Otto S, Ravi D, et al. Cadherin-9 regulates synapse-specific differentiation in the developing hippocampus. Neuron 2011, 71: 640–655.

[69] Deciphering Developmental Disorders S. Prevalence and architecture of de novo mutations in developmental disorders. Nature 2017, 542: 433–438.

[70] Wang K, Zhang H, Ma D, Bucan M, Glessner JT, Abrahams BS, et al. Common genetic variants on 5p14.1 associate with autism spectrum disorders. Nature 2009, 459: 528–533.

[71] Kerin T, Ramanathan A, Rivas K, Grepo N, Coetzee GA, Campbell DB. A Noncoding RNA Antisense to Moesin at 5p14.1 in Autism. Science Translational Medicine 2012, 4(128): 128ra40.

[72] DeWitt JJ, Grepo N, Wilkinson B, Evgrafov OV, Knowles JA, Campbell DB. Impact of the Autism-Associated Long Noncoding RNA MSNP1AS on Neuronal Architecture and Gene Expression in Human Neural Progenitor Cells. Genes (Basel) 2016, 7(10): 76.

[73] DeWitt JJ, Hecht PM, Grepo N, Wilkinson B, Evgrafov OV, Morris KV, et al. Transcriptional Gene Silencing of the Autism-Associated Long Noncoding RNA MSNP1AS in Human Neural Progenitor Cells. Dev Neurosci 2016, 38: 375–383.

[74] Grove J, Ripke S, Als TD, Mattheisen M, Walters RK, Won H, et al. Identification of common genetic risk variants for autism spectrum disorder. Nat Genet 2019, 51: 431–444.

