## Supplementary figures and images for "Association of *CDH11* with ASD revealed by matched-gene co-expression analysis and mouse behavioral studies"

### Figure_S1

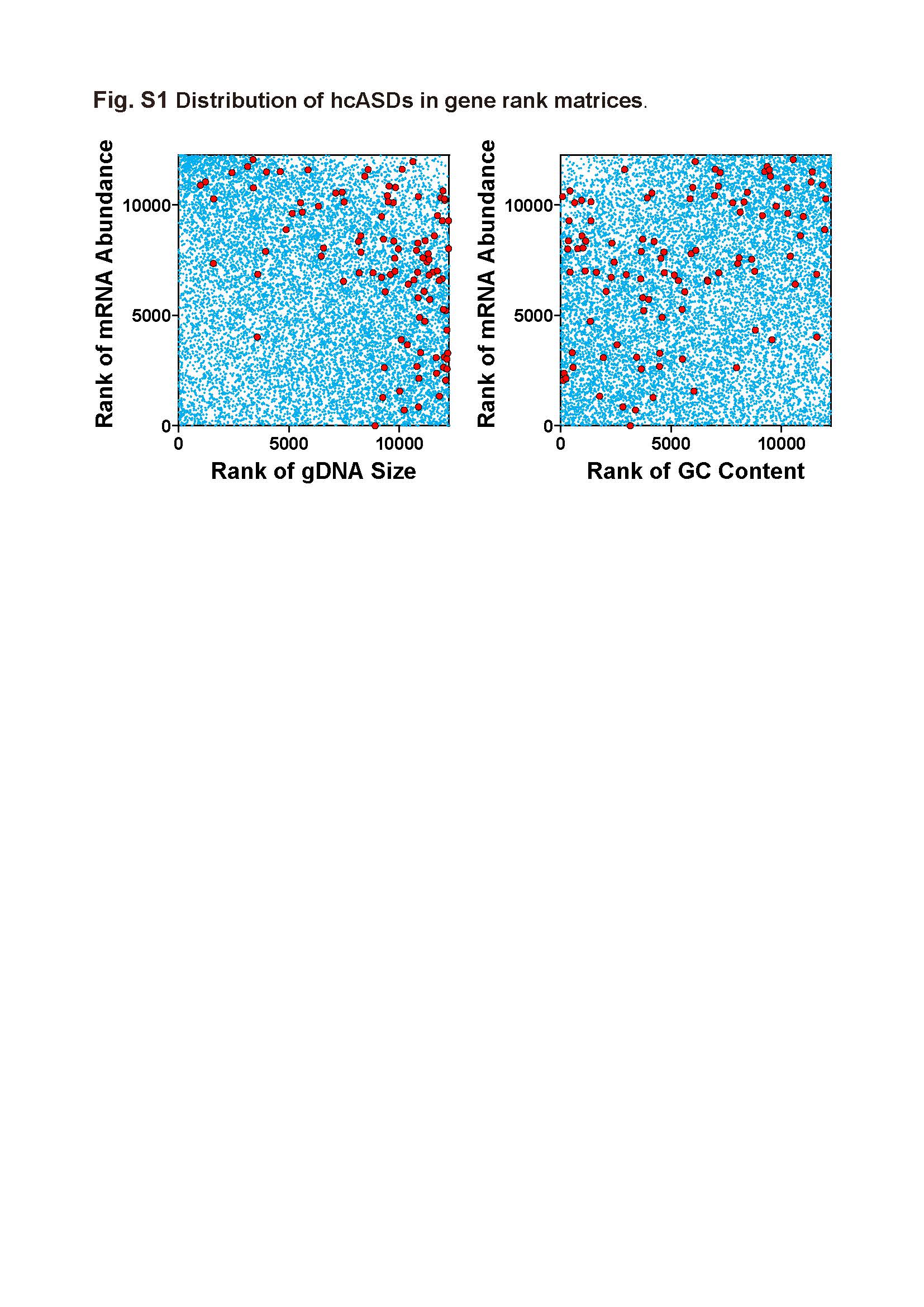

### Figure_S2

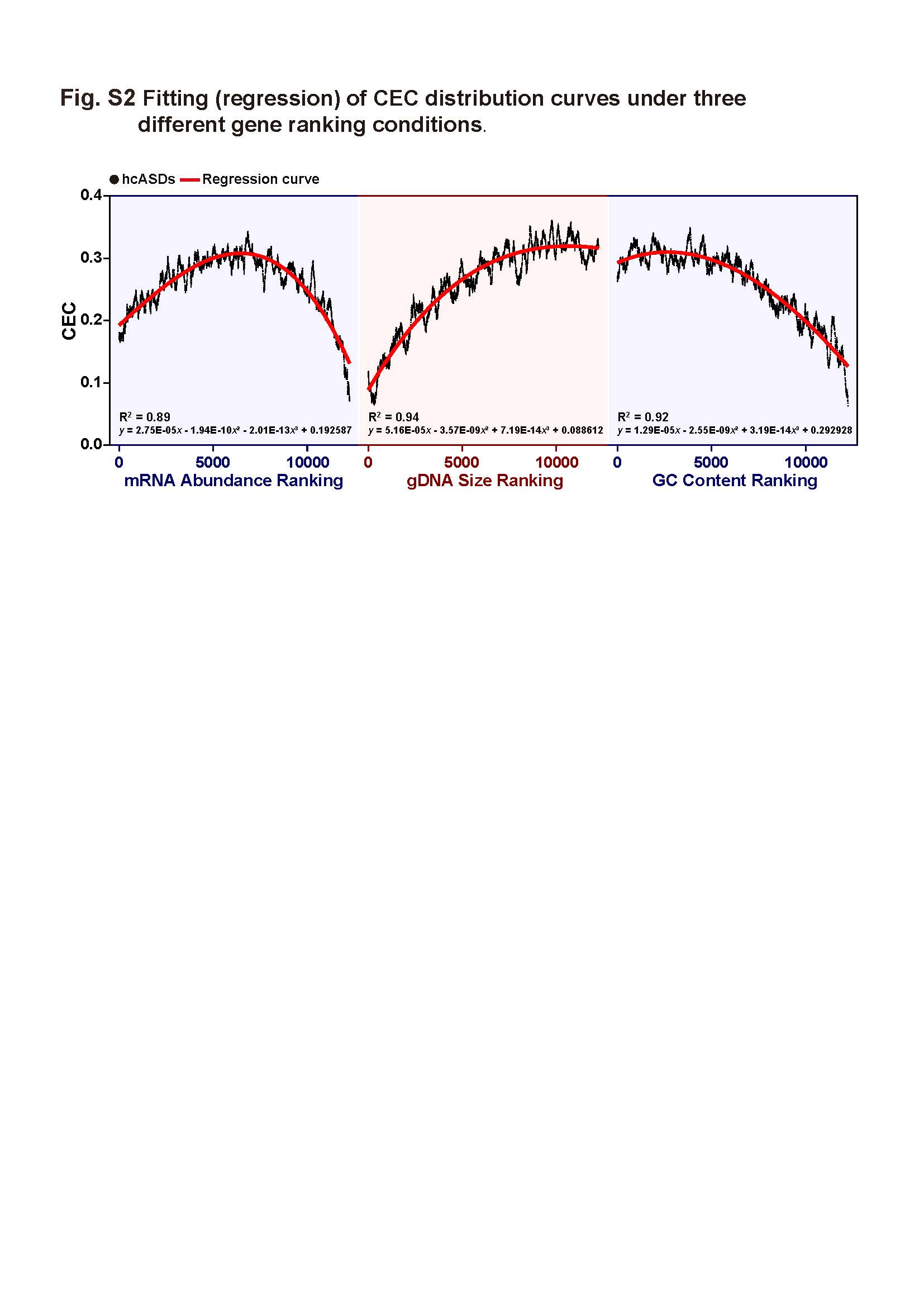

### Figure_S3

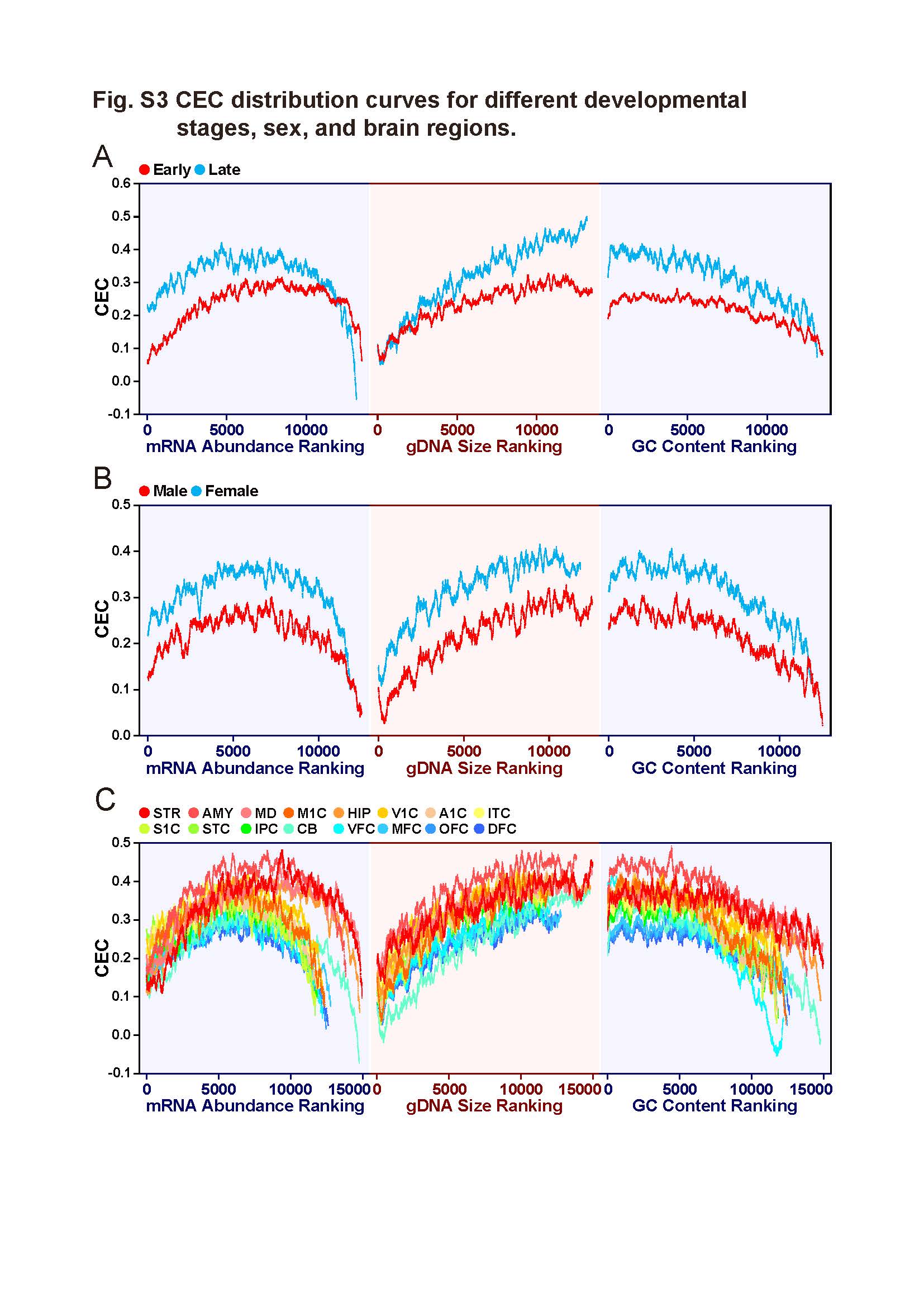

### Figure_S4

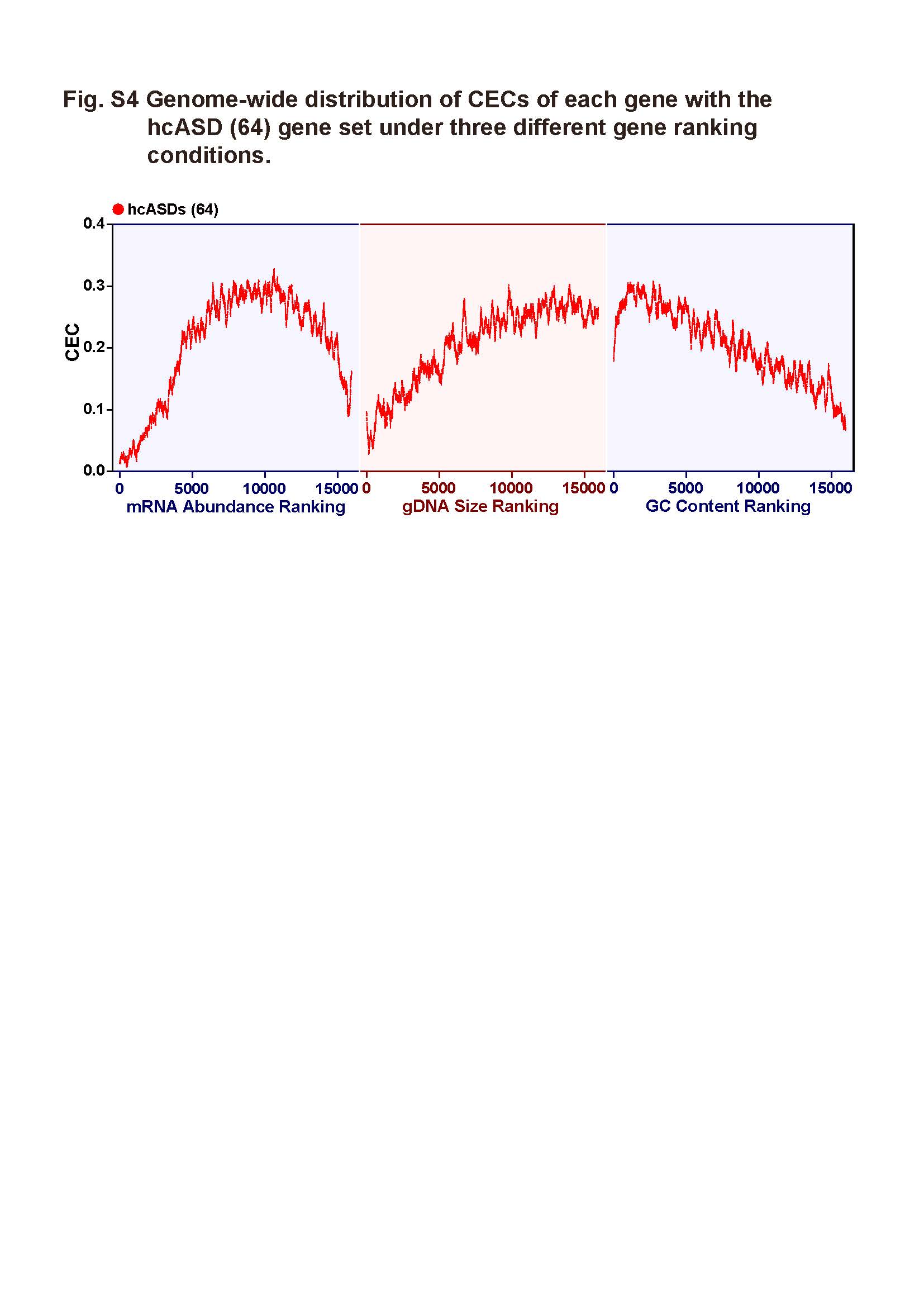

### Figure_S5

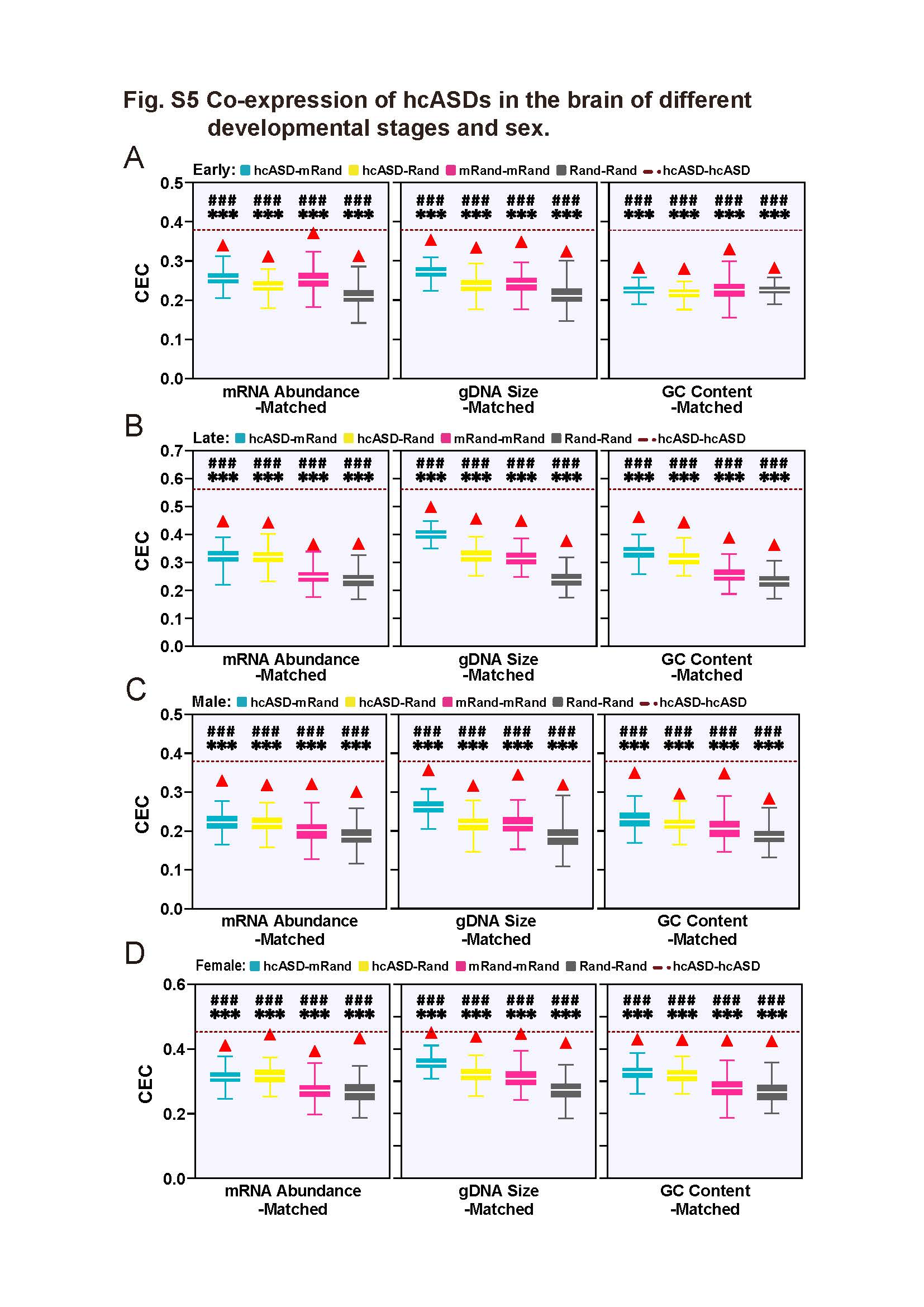

### Figure_S6

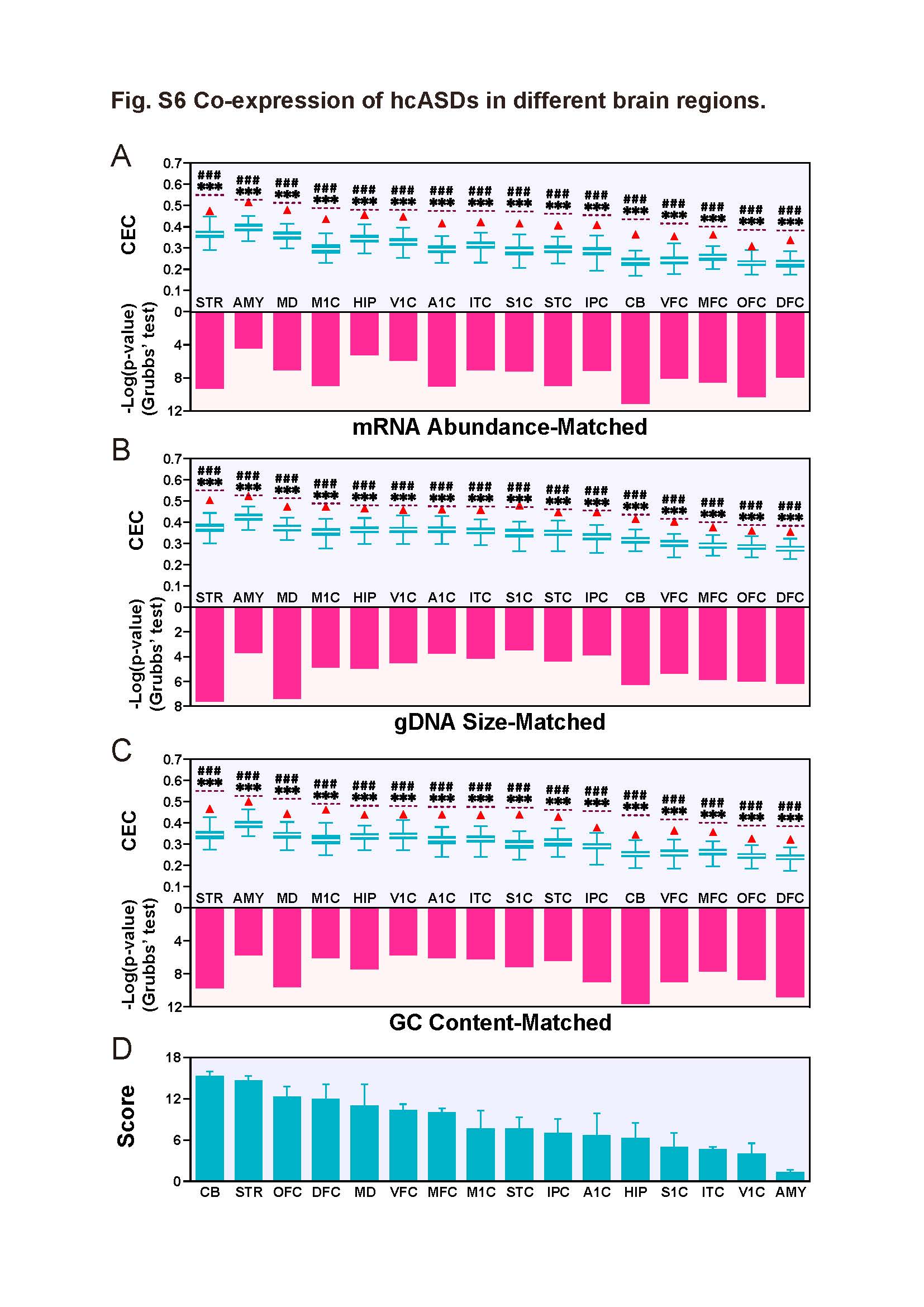

### Figure_S7

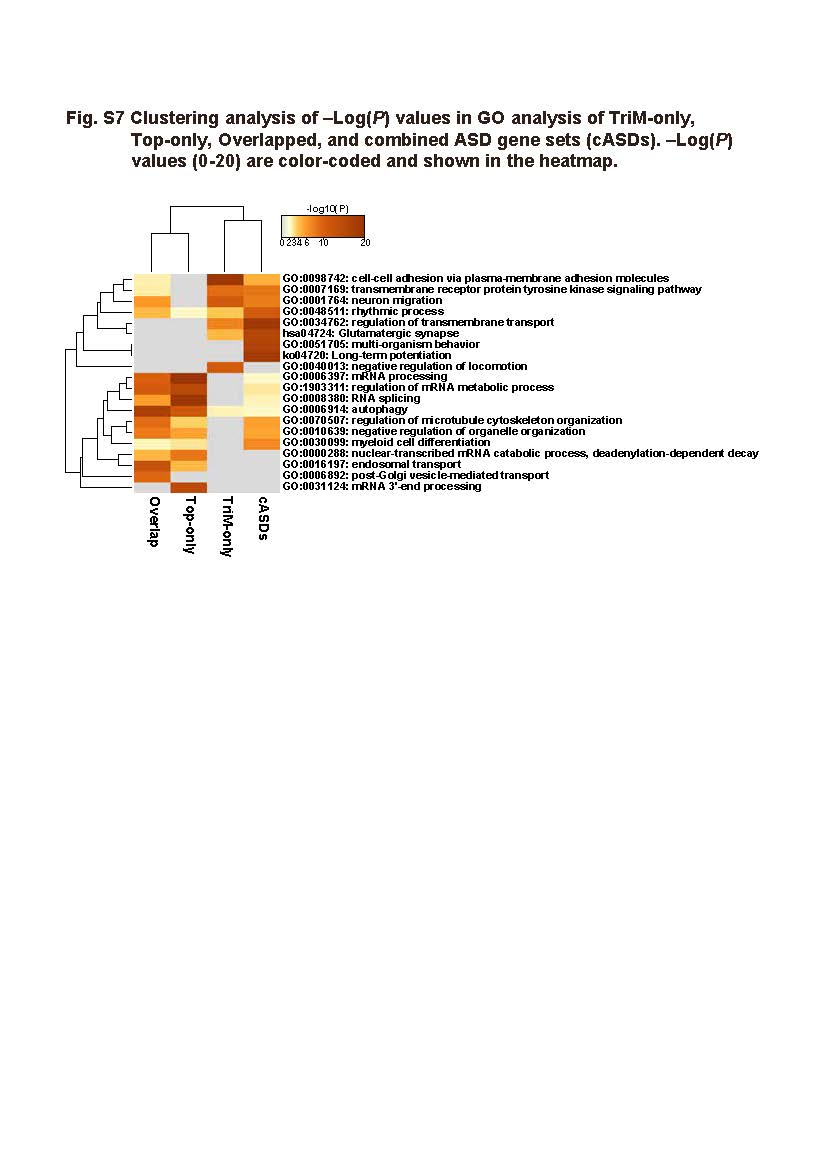
